# Genetic effects on brain traits impact cell-type specific gene regulation during neurogenesis

**DOI:** 10.1101/2020.10.21.349019

**Authors:** Nil Aygün, Angela L. Elwell, Dan Liang, Michael J. Lafferty, Kerry E. Cheek, Kenan P. Courtney, Jessica Mory, Ellie Hadden-Ford, Oleh Krupa, Luis de la Torre-Ubieta, Daniel H. Geschwind, Michael I. Love, Jason L. Stein

## Abstract

Interpretation of the function of non-coding risk loci for neuropsychiatric disorders and brain-relevant traits via gene expression and alternative splicing is generally performed in bulk post-mortem adult tissue. However, genetic risk loci are enriched in regulatory elements active during neocortical differentiation, and regulatory effects of risk variants may be masked by heterogeneity in bulk tissue. Here, we map expression quantitative trait loci (eQTLs), splicing QTLs (sQTLs), and allele specific expression in primary human neural progenitors (n=85) and their sorted neuronal progeny (n=74), identifying numerous loci not detected in either bulk developing cortical wall or adult cortex. Using colocalization and genetic imputation via transcriptome wide association, we uncover cell-type specific regulatory mechanisms underlying risk for brain-relevant traits that are active during neocortical differentiation. Specifically, we identified a progenitor-specific eQTL for *CENPW* co-localized with common variant associations of cortical area and educational attainment.

## Introduction

Genome wide association studies (GWAS) have identified many common non-coding variants associated with risk for brain structure, neurodevelopmental disorders, and other brain-related traits ^1–7^. However, it is challenging to determine the mechanism of non-coding variants because, in general, (1) the genes impacted by non-coding risk variants are unknown, (2) the cell type(s) and developmental period(s) where the variants have an effect are not known, (3) and there may be limited availability of tissue representing the causal developmental stage and cell type.

One potential mechanism by which non-coding genetic variation can influence brain traits is through alterations in gene expression, or expression quantitative trait loci (eQTLs). Genetic variation also impacts transcript splicing ^8–10^, and several studies have implicated genetically mediated alterations in splicing as important risk factors for neuropsychiatric disorders ^11–13^.

Most current efforts to explain the function of these risk loci rely on mapping local expression and splicing quantitative trait loci (e/sQTLs) in bulk adult brain tissue ^14, 15^. However, genetic risk loci are enriched in cell types relevant for neocortical differentiation that are not present in the adult brain ^16, 17^. e/sQTL studies have been performed on human fetal brain bulk cortical tissue ^18–20^, but there are advantages to a cell-type specific approach given that regulatory effects of risk variants may be masked by heterogeneity in bulk tissue ^21–23^.

Utilizing a cell-type specific *in vitro* model system including neural progenitors (N_donor_ = 85) and their virally labelled and sorted neuronal progeny (N_donor_ = 74) derived from a multi-ancestry population, here we investigated how common genetic variants impact brain related traits through gene expression and splicing during human neurogenesis. We discovered 2,079/872 eQTLs in progenitors and neurons, and 5,676/4,590 sQTLs in progenitors and neurons, respectively. Importantly, 66.1%/47% of eQTLs in progenitor/neuron and 78.8%/76.1% sQTLs in progenitor/neuron were unique, and were not found in fetal bulk brain e/sQTLs from a largely overlapping sample ^19^ or in adult bulk e/sQTL data from GTEx project ^24^. We showed both eQTLs and sQTLs colocalized with known GWAS loci for neuropsychiatric disorders and other brain relevant traits in a cell-type specific manner. By integrating the dataset generated here with cell-type specific chromatin accessibility from the same cell lines ^16^ and brain structure GWAS ^4^, we propose a regulatory mechanism whereby genetic variation influences a proxy of human intelligence across multiple levels of biology. Furthermore, we genetically imputed disease/brain trait susceptibility gene expression and alternative splicing in these cell types using transcriptome-wide association study (TWAS), that identified cell type and temporal specific risk genes and introns.

## Results

### Transcriptomic profiles of primary human progenitors and neurons recapitulates cell-type specific characteristics of cortical development

We established an *in vitro* culture of primary human neural progenitor cell (phNPC) lines derived from genotyped human fetal brain tissue (N = 89 unique donors) at 12-19 post conceptional weeks (PCW) (14-21 gestation weeks), that recapitulates the developing human neocortex ^25–28^ (Figure 1A, Methods). Immunofluorescence of the cells showed that undifferentiated progenitors expressed PAX6 and SOX2 (90-95%), consistent with a homogenous culture of radial glia ^29, 30^ (Figure 1B). At 5 weeks post-differentiation, phNPC cultures were transduced with a virus which expresses EGFP in neurons (AAV2-hSyn1-EGFP), which enabled us to isolate neurons via FACS sorting at 8 weeks post-differentiation (Figure 1A and 1B, Figure S1A and S1B, Methods).

**Figure 1.**
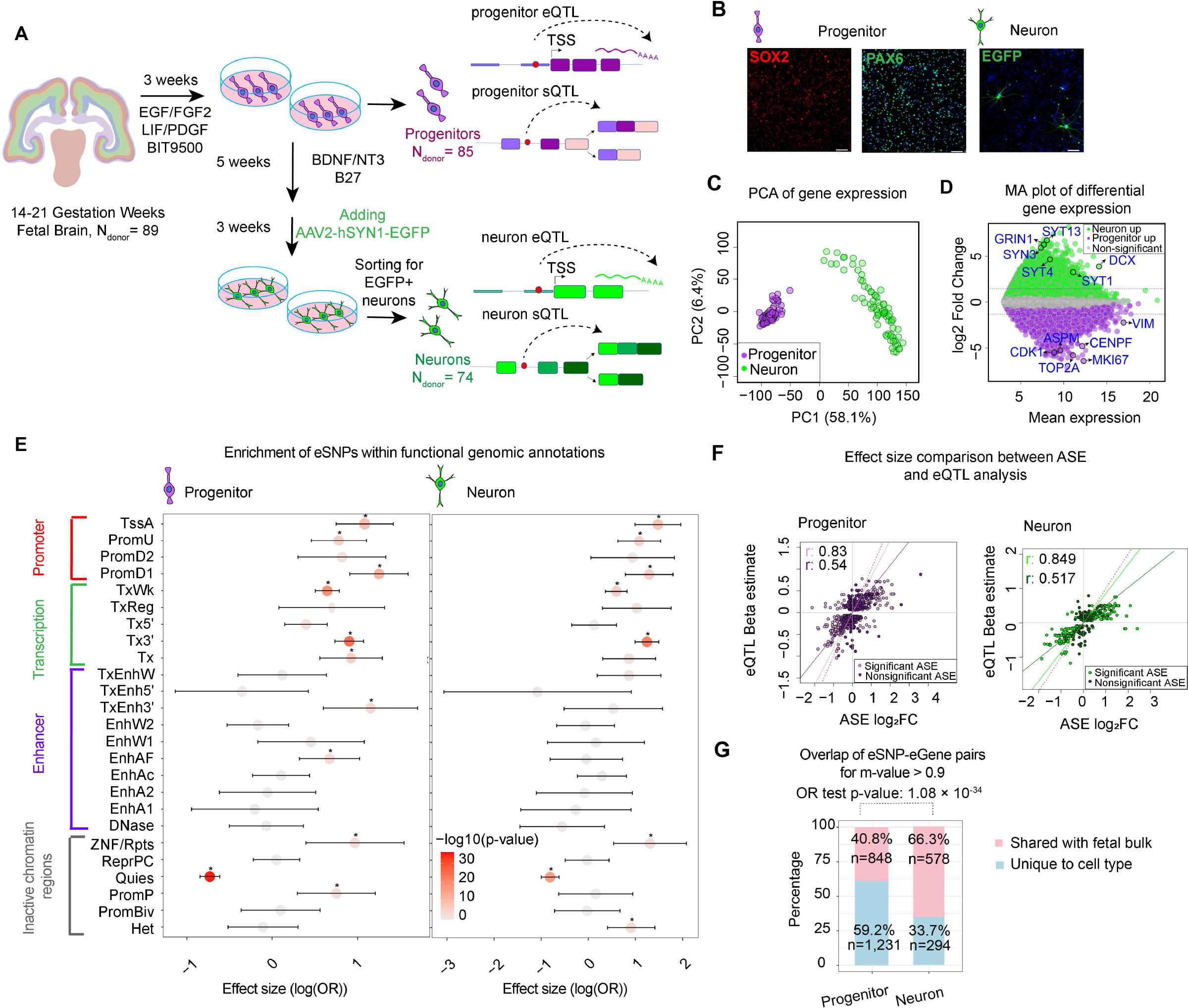
Study design and cell-type specific eQTL analysis. **(A)** Study design illustrating the fetal brain tissue derived cell-type specific system to perform eQTL and sQTL analysis. **(B)** Immunofluorescence of the cells showed that undifferentiated progenitors expressed SOX2 (in red) and PAX6 (in green), and expression of EGFP (in green) in 8-week differentiated neurons labelled with AAV2-hSyn1-EGFP (scale bar is 100 μm, DAPI in blue). **(C)** Principal component analysis of progenitor (purple) and neuron (green) transcriptomes from each donor indicates cell-type specific clustering. **(D)** MA plot showing differentially expressed genes in progenitor versus neurons. log2FC > 0 and adjusted p-value < 0.05 indicates genes upregulated in neurons shown in green (Neuron up), log2FC < 0 and adjusted p-value < 0.05 indicates genes upregulated in progenitors shown in purple (Progenitor up) and genes not significantly differentially expressed between two cell types are shown in grey. Blue lines indicates |log_2_FC| > 1.5. **(E)** Enrichment of progenitor eSNPs (left), and neuron eSNPs (right) within chromatin states in the fetal brain from chromHMM listed on the y-axis. The x-axis shows the effect size of enrichment with 95% upper and lower confidence interval and the plot is color-coded based on −log10(p-value) value from enrichment analysis. Significant enrichments are shown with an asterisk. Enrichment was tested using eQTLs thresholded at the eigenMT-BH p-value. **(F)** Comparison of the effects of shared ASE sites and eQTLs in progenitors (left in purple) and neurons (right in green). Nonsignificant ASE sites are shown as darker colors for both cell types, and significant ASE sites are shown as lighter colors. Correlation coefficient (r) values are indicated in colors for each category and the red dashed line indicates x = y. **(G)** Overlap percentage of cell-type specific eSNP-eGene pairs shared with fetal bulk eQTLs, respectively at m-value > 0.9. Odds ratio (OR) test p-values are shown.

We acquired transcriptomic profiles of progenitors and neurons via RNA-sequencing, observing a strong correlation of libraries from the same donor (Figure S1C). After correction for technical confounds (Figure S1D), progenitors and neurons clustered separately by principal component analysis (PCA) of global gene expression, indicating global transcriptomic differences by cell type (Figure 1C). Both cell types showed expression of a variety of known cell-type specific markers in the literature (Figure S1E). Next, we identified differentially expressed genes, which were involved in cell cycle and neurotransmission upregulated in progenitors and neurons, respectively (Figure 1D and Figure S2F, Table S1).

We evaluated how well the *in vitro* progenitors and neurons we generated model in vivo neurodevelopment. We implemented the transition mapping (TMAP) approach for a global assessment of transcriptomic overlap between *in vitro* cultures and *in vivo* post-mortem human brain samples, as described in our previous work ^26^ (Methods). We compared the transition from progenitor to neurons with laser capture microdissection of cortical laminae from postmortem human fetal brain at 15-21 PCW ^31^. We observed the strongest overlap in the transition from progenitors to neurons with the transition from outer subventricular zone (oSVZ) to intermediate zone (IZ) or subplate zone (SP) (Figure S1G), supporting the *in vivo* fidelity of our culture system representing neurogenesis during mid-fetal development.

### Discovery of cell-type specific genetically altered gene expression via local expression quantitative loci (eQTL) analysis

To investigate the impact of genetic variation on gene expression, we performed a local eQTL analysis by testing the association of each gene’s expression levels with genetic variants residing within ±1 Mb window of its transcription start site (TSS) ^32, 33^ (Figure S2A, see Methods). We implemented a linear mixed effects model to stringently control for population stratification using a kinship matrix as a random effect with inferred technical confounders as fixed effects, separately for each cell type (λ_GC_ for progenitor = 1.028 and λ_GC_ for neuron = 1.007; see Methods, Figure S2B-S2G). After retaining associations which were lower than 5% false discovery rate with a hierarchical multiple testing correction ^34, 35^ (Methods), we obtained conditionally independent eQTLs (Figure S2B and S2C, see Methods). We identified 1,741 eGenes with 2,079 eSNP-eGene pairs in progenitors and 840 eGenes with 872 eGene-eSNP pairs in neurons (Figure S3A and Table S1). 47%/68% of progenitor/neuron conditionally independent eQTLs were shared with m-value > 0.9 (Figure S3B).

We determined if eSNPs were enriched in chromatin states of the fetal human brain ^17, 36^ (Figure 1E). Both progenitor and neuron specific eSNPs were enriched in promoters, enhancers and actively transcribed sites present in the fetal brain, but depleted mostly in quiescent chromatin regions. Importantly, 32.7%/34.3% of progenitor/neuron specific eQTLs, respectively, were supported by cell-type specific allele specific expression (ASE), that is not subject to cross donor technical confounding, like population stratification ^33, 37, 38^ (Figure S3E-S3H, Table S2). These shared eQTLs with genome-wide significant ASE sites were highly concordant in effect size and direction (Figure 1F).

### Cell-type specific eQTLs exhibits both cell-type and temporal specificity

We aimed to determine the utility of our cell-type specific eQTL study by comparison to pre-existing bulk brain eQTL studies. Comparing our results to a bulk fetal cortical wall eQTL dataset from a previous study using a partially overlapping set of donors ^19^, we observed that 40.8%/66.3% of progenitor/neuron eQTLs were also detected in the fetal bulk eQTLs (Figure 1G, m-value > 0.9 indicates shared effects; odds ratio test between cell type sharing with fetal bulk: p-value: 1.08 × 10^-34^, see Figure S3C for LD-based overlap). Taken together, our observations propose that both cell-type specific eQTLs, but especially progenitor eQTLs in our cell-type specific system offer novel regulatory mechanisms which can provide additional information beyond existing prenatal datasets ^18–20^.

We next explored the temporal specificity of cell-type specific eQTLs by utilizing adult brain bulk cortical eQTL data from the GTEx project ^24^. We observed 18.9%/28.3% of eSNP-eGene pairs in progenitors and neurons, respectively, were also found in adult brain eQTL data (Figure S3D). That suggests largely independent genetic mechanisms regulating genes from development to adulthood ^20^.

### Cell-type specific identification of splicing quantitative trait loci (sQTL) in developing human brain

Given the impact of genetic variation on alternative splicing ^9, 11, 19, 39^, we next performed a splicing quantitative loci (sQTL) analysis separately within progenitors and neurons. We quantified alternative intron excisions as percent spliced in (PSI) by implementing the Leafcutter software, an annotation free approach that allows for discovery of novel isoforms ^40^. We found 34,449/35,285 intron excisions present more often in progenitors/neurons, respectively (see Methods, Table S3). As a specific example, we found a differential alternative splicing site within the *DLG4* gene encoding the postsynaptic density protein 95 (PSD-95). An exon skipping splice site supporting nonsense mediated decay (splice 1, ENST00000491753) was upregulated in progenitors; while another splice site supporting multiple protein coding transcripts (splice 2) was upregulated in neurons (Figure 2A). Post-transcriptional repression of PSD-95 expression in neural progenitors via nonsense mediated decay at splice 1 site has been previously experimentally validated ^41, 42^, giving strong confidence in the cell-type specific splicing calls.

**Figure 2.**
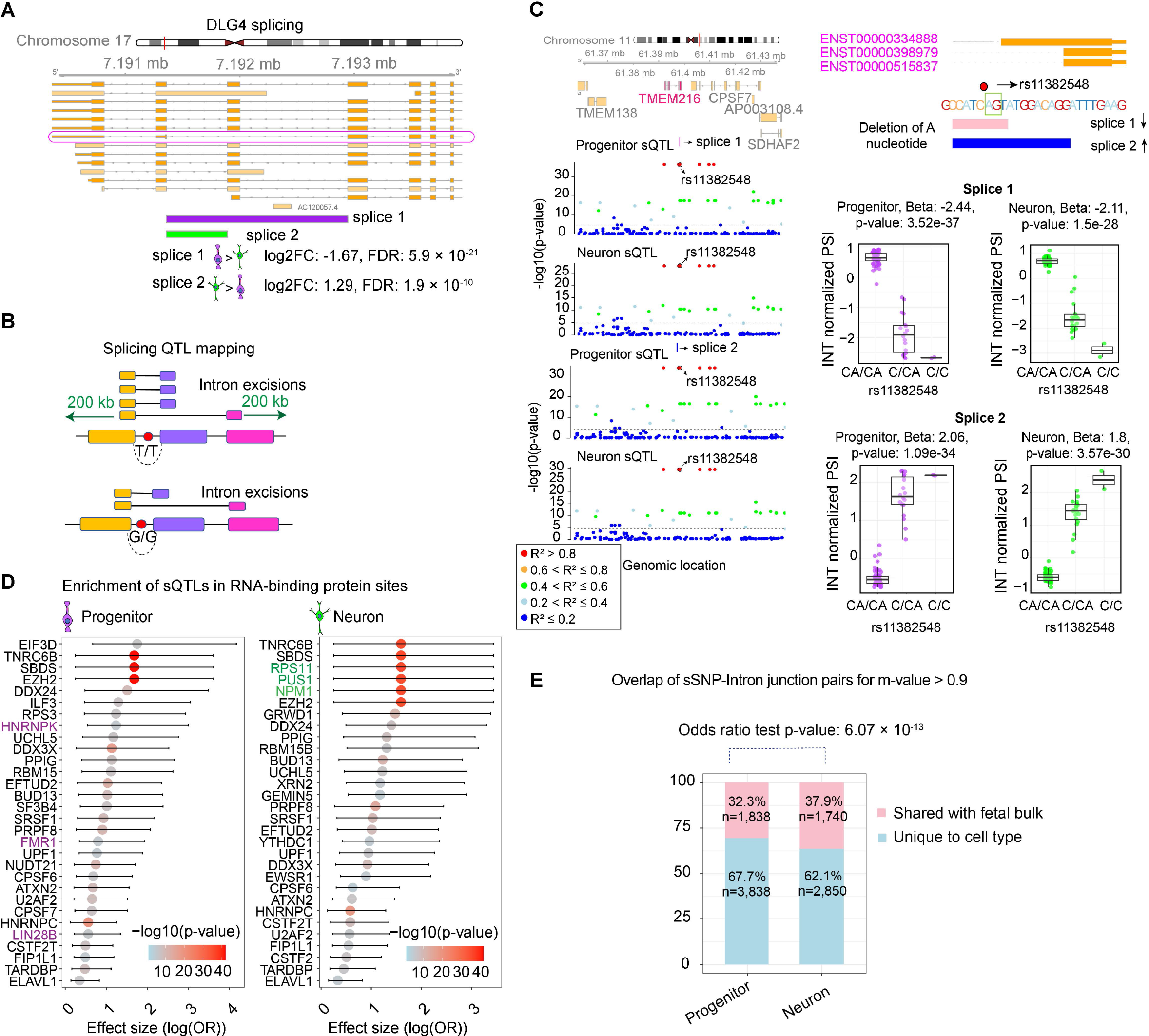
Cell-type specific sQTL analysis. **(A)** Differential splicing of two intron junctions within the *DLG4* gene. Splice 1 (chr17:7191358-7192945) supports a previously validated nonsense mediated decay transcript (ENST00000491753) with higher expression in progenitors, whereas splice 2 (chr17:7191358-7191893) has higher expression in neurons. **(B)** A schematic illustrating splicing QTL mapping. Association of variants locating within 200 kb distance from each end of intron junctions were tested. The T allele is associated with more frequently splicing of the shorter intron junction. **(C)** Two intron junctions supporting an alternative 3’ splicing site for *TMEM216* gene regulated by variant rs11382548 located at the splice site. The regional association of variants to two introns is shown in the genomic track at the left colored by pairwise LD r^2^ relative to variant rs11382548, association p-values on the y-axis, and genomic location of each variant on the x-axis. Gene model of *TMEM216* is shown in the upper right with position of the variant rs11382548, green box indicates splice site. Box plots in the lower right show quantile normalized PSI values for splice 1 (chr11:61397975-61398261) and splice 2 (chr11:61397975-61398270) across variant rs11382548. **(D)** Enrichment of cell-type specific sQTLs within RNA-binding protein (RBP) binding sites based on a CLIP-seq dataset. The top 30 RBPs based on −log10(enrichment p-value) are listed on the y-axis, and the x-axis shows the effect size from enrichment test, where data points colored by −log10(p-value) from the enrichment test and cell-type specific RBPs are colored with purple for progenitors at the left, and as green for neuron at the right. **(E)** Overlap percentage of cell-type specific sSNP-intron junction pairs shared with fetal bulk sQTLs, respectively at m-value > 0.9. Odds ratio (OR) test p-values are shown.

For the sQTL analysis, we implemented an association test between PSI of each intron excision and genetic variants located within a ± 200 kb window from the start and end of the splice junctions (Figure 1A and 2B). We retained significant associations which were lower than 5% false discovery rate by implementing a hierarchical multiple testing correction (see Methods), and applied conditional analysis to identify independent sQTLs (Figure S2C, S2E and S2H). We identified 4,708 intron excisions associated with 5,676 sSNPs-intron junction pairs in progenitors and 4,039 intron excisions associated with 4,590 sSNPs-intron junction in neurons (Figure S4A and S4B, 52.9%52.3% of progenitor/neuron conditionally independent sQTLs were shared with m-value > 0.9, Table S3). As an example, we found that the indel variant rs11382548 creating a canonical splice acceptor sequence impacted two different intron excisions supporting alternative 3’ splice sites for *TMEM216* gene (Figure 2C). Deletion of the A nucleotide at a canonical splice acceptor site of the last exon of *TMEM216* leads to disruption of the alternative splicing event for transcript ENST00000334888, and increased usage of transcript ENST00000398979 and ENST00000515837 in both progenitors and neurons. This sQTL may be relevant to neurogenesis because knockdown of the *TMEM216* gene reduces division of both apical and intermediate progenitor cells during corticogenesis ^43^.

Interestingly, many splice sites were previously unannotated in the gene models we used (Ensembl Release 92). We detected 8.8%/11.3% cryptic at the 5’ end; 12%/12% cryptic 3’end; 8.9%/10.6% both cryptic ends for intron excision within progenitors/neurons.

We also found that cell-type specific sQTLs in progenitors/neurons were enriched for RNA binding sites from CLIP-seq databases of 74/76 different RNA-binding proteins (RBP) ^44^ (Figure 2D, Table S3). Strikingly, 22/24 of those proteins were specifically enriched in progenitor and neuron specific sQTLs, respectively. Among RBP binding sites specifically enriched for progenitor sQTLs, we found LIN28B, known to play a role in neural progenitor proliferation and differentiation ^45^. On the other hand, for neurons, we detected enrichment of the NPM1 regulating neuronal survival ^46^. These observations suggest that sQTLs alter the function of RBPs with cell-type specific splicing roles during neural development.

In order to examine if variants associated with alternative splicing also alter expression of the same genes, we compared cell-type specific sQTLs with cell-type specific eQTLs. Only 17.6%/9.5% of sGenes, the genes that harbor intron excisions, were also eGenes for progenitors and neurons eQTLs, respectively (Figure S4C, upper panel). Furthermore, we also found that only 2.9% and 1.4% of sGene-sSNP pairs overlapped with eGene-eSNP pairs for progenitors and neurons, respectively (Figure S4C, lower panel), indicating that sQTLs generally function through independent mechanism from eQTLs.

We next examined the impact of cell-type specificity on identification of sQTLs. 32.3%/37.9% of progenitor/neuron sQTLs were also detected in the fetal bulk sQTLs (Figure 2E, m-value > 0.9 indicates shared effects; odds ratio test between cell type sharing with fetal bulk: p-value: 6.07 × 10^-13^, see Figure S4D for LD-based overlap). A smaller overlap of progenitor sQTLs with bulk cortical fetal tissue as compared to neuron sQTLs indicated that cell-type specificity allowed for novel discovery of progenitor sQTLs. Also, we found 6%/2.4% of sSNP-intron junction pairs in progenitors and neurons, respectively shared with adult brain bulk cortical sQTL data from GTEx project ^24^ (Figure S4E), showing temporal specificity of cell-type specific sQTLs.

### Cell-type specific e/sQTLs proposes regulatory mechanisms of brain related GWAS

We sought to explain the regulatory mechanism of individual loci associated with neuropsychiatric disorders, brain structure traits, and other brain-relevant traits by leveraging genetic variants regulating cell-type specific gene expression and splicing. We co-localized GWAS loci of these traits with cell-type specific eQTLs and sQTLs using a conditional analysis to ensure the loci were shared across traits ^47^ (see Methods for the list of GWAS used for this analysis).

We discovered 41, 13, and 20 GWAS loci that co-localized specifically with progenitor eQTL, specifically with neuron eQTLs, or with both cell types, respectively (Figure 3A, Table S4). These observations show that the same genetic variants impact gene expression, neuropsychiatric traits, and brain structure in a cell-type specific manner. Importantly, 98 trait associated loci-gene pairs (one locus could be associated with multiple different genes) were not found using fetal bulk cortical tissue eQTLs, where tissue heterogeneity may have masked their detection (Figure 3B).

**Figure 3.**
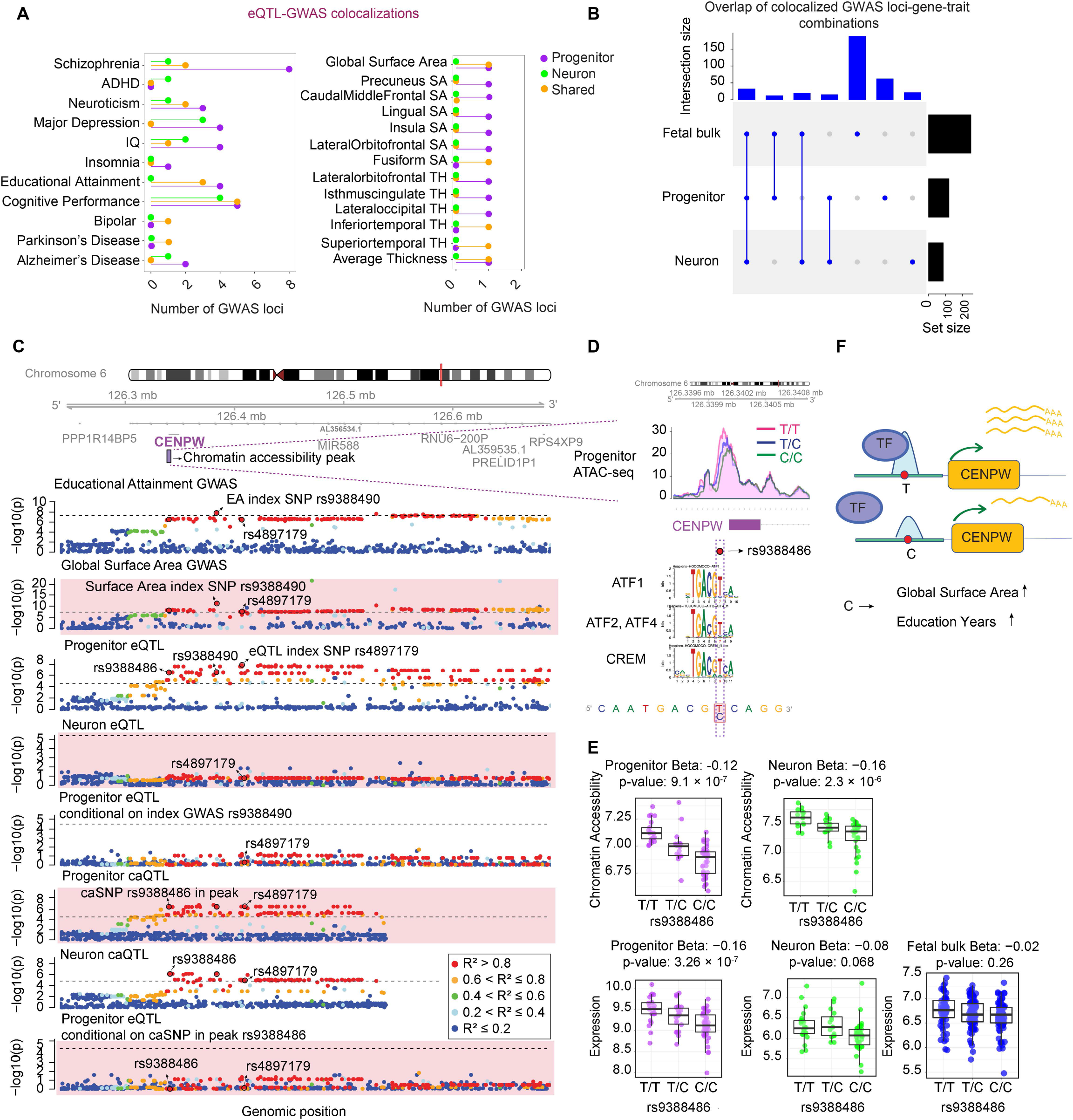
Colocalization of cell-type specific eQTLs with GWAS for brain related traits. **(A)** Number of GWAS loci colocalized with progenitor (purple), neuron (green) specific eQTLs or both cell types (orange). Each GWAS trait is listed on the y-axis (SA: surface area, TH: thickness). **(B)** Overlap of colocalized GWAS loci-gene pairs per trait combinations across progenitor, neuron and fetal bulk eQTL colocalizations for the traits listed in Figure 3A. **(C)** Genomic track showing regional association of variants with educational attainment (EA), global surface area (GSA) and *CENPW* gene expression in progenitors and neurons, association p-values (p) on the y-axis, and genomic location of each variant on the x-axis. Progenitor eSNP rs4897179 (3rd row) was coincident with index SNP (rs9388490) for both EA (1st row) and GSA GWAS (2nd row), and conditioning progenitor eSNP rs4897179 on rs9388490 showed colocalization of the two signals (5th row). Also, rs4897179 was colocalized with another variant (rs9388486) located in the chromatin accessibility peak at the promoter of *CENPW* gene (6th and 8th rows). Genomic tracks were color-coded based on LD r^2^ relative to the variant rs9388486. **(D)** Plot showing the chromatin accessibility peak (chr6:126339531-126340960) in progenitors across different genotypes of rs938848. The C allele of rs9388486 disrupted binding motifs of transcription factors including CREM, ATF1, ATF2 and ATF4. **(E)** Box plots showing chromatin accessibility across rs9388486 genotypes in progenitors (purple) and neurons (green) (upper panel). Box plots showing VST normalized *CENPW* gene expression across rs9388486 genotypes in progenitors (purple), neurons (green) and fetal bulk (blue) (bottom panel). **(F)** A schematic showing that the implicated transcription factor has decreased preference to bind at the C allele, which results in lower *CENPW* expression, increase in global surface area and educational attainment.

Next, we leveraged our cell-type specific chromatin accessibility QTL (caQTL) dataset ^16^ together with eQTLs in order to explain the regulatory mechanism underlying GWAS loci associated with brain relevant traits. As a specific example, we found a colocalization of a locus within the *CENPW* gene across caQTLs, eQTLs, GWAS for Global Surface Area (GSA) and for Educational Attainment (EA) (Figure 3C). The progenitor index eSNP rs4897179 that was not detected in bulk cortical fetal tissue eQTLs (nominal p-value = 3.26 × 10^-7^ in progenitors, nominal p-value = 0.068 in neurons, and nominal p-value = 0.26 in fetal cortical bulk tissue), for the *CENPW* eGene, was colocalized with variant rs9388490, which is the index SNP for both GSA and EA GWAS (nominal p-value = 4.95 × 10^-12^ in GSA GWAS, and nominal p-value = 1.43 × 10^-8^ in EA GWAS). Also, we found that a SNP (rs9388486) located within a chromatin accessible peak region 107 bp upstream of TSS of the *CENPW* gene was colocalized with the index eSNP. We therefore consider rs9388486 as the potential causal variant, and noted that the C allele disrupts the motifs of the transcription factors CREM, ATF2, ATF4 and ATF1 (Figure 3D). CENPW is required for appropriate kinetochore formation and centriole splitting during mitosis ^48^, and increased CENPW levels lead to apoptosis in the developing zebrafish central nervous system ^49^. Overall, these observations propose a cell-type specific mechanism whereby the C allele at variant rs9388486 disrupts transcription factor binding and diminishes accessibility at the *CENPW* gene promoter, resulting in decreased *CENPW* gene expression levels in progenitors (Figure 3E and 3F), presumably altering neurogenesis or reducing apoptosis, leading to increased cortical surface area and higher cognitive function.

We also aimed to examine cell-type specific splicing QTLs colocalized with GWAS loci. We observed 28, 23, and 29 GWAS loci in total that co-localized with specifically progenitor/neuron sQTLs and sQTLs present in both cell types (Figure 4A, Table S4). Similar to eQTL colocalizations, we observed that 124 trait associated loci-intron junction pairs were detected only with cell-type specific sQTL (one locus could be associated with multiple intron junctions), but not fetal bulk cortical sQTLs (Figure 4B). Interestingly, we detected a progenitor sSNP (rs3740400, nominal p-value: 3.29 × 10^-10^) associated with an unannotated exon skipping splicing event for the *AS3MT*, that was not detected in fetal bulk sQTL data (nominal p-value: 0.005), was colocalized with an index GWAS SNP for schizophrenia (rs11191419, ^1^ (Figure 4C). The risk allele for schizophrenia (T) was associated with more usage of this splicing site (Figure 4D and 4E). A transcript supported by the same exon skipping event was discovered previously in the adult brain, that was regulated by variant rs7085104 in LD with rs3740400 ^50^. Here, we demonstrate genetically regulated upregulation of this transcript in neural progenitors, but not early born neurons, which shows a novel developmental basis for this previously identified association. We also observed that another progenitor specific sSNP (rs1222218) regulating a novel alternative exon skipping event for *ARL14EP* gene, was colocalized with SCZ index SNP (rs1765142) ^1^ (Figure 4F). The risk allele for SCZ led to more frequent skipping of the exon, supporting expression of a novel isoform (Figure 4G and 4H). *ARL14EP* gene has been shown to play a role in axonal development in the mouse neurons ^51^. Here, we propose a novel transcript of this gene with expression in progenitors as a risk factor for SCZ.

**Figure 4.**
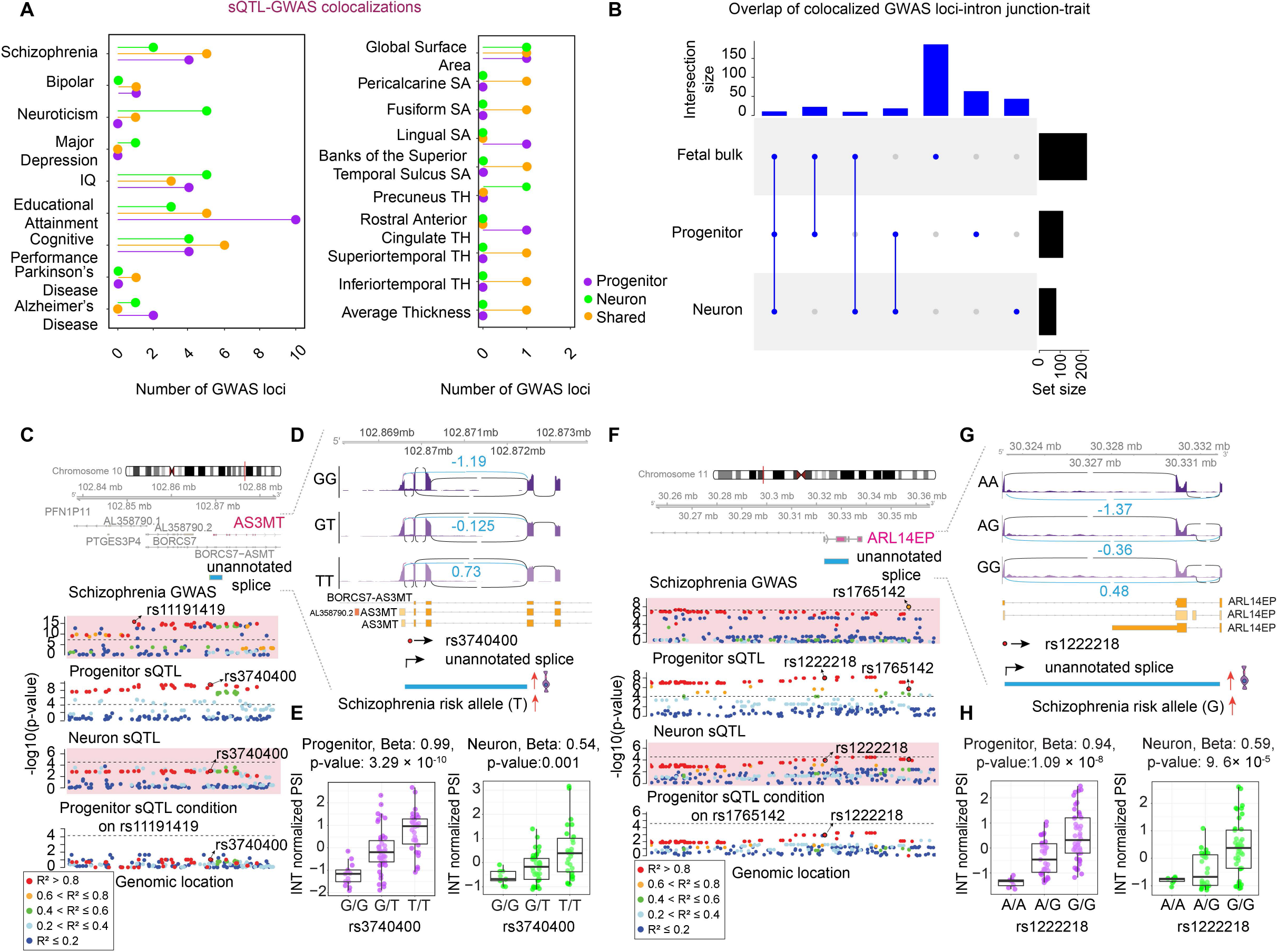
Colocalization of cell-type specific sQTLs with GWAS for brain related traits. **(A)** Number of GWAS loci colocalized with progenitor (purple), neuron (green) specific sQTLs or both cell types (orange). Each GWAS trait is listed on the y-axis (SA: surface area, TH: thickness). **(B)** Overlap of colocalized GWAS loci-intron junction pairs per trait across progenitor, neuron and fetal bulk sQTL colocalizations for the traits listed in Figure 4A. **(C)** Genomic tracks showing regional association of variants with SCZ and an unannotated exon skipping splice site (chr10:102869593-102872448) for *AS3MT* gene in progenitors and neurons association p-values on the y-axis, and genomic location of each variant on the x-axis. The splice site was associated with progenitor sSNP (rs3740400) colocalized with SCZ GWAS index SNP (rs11191419). Genomic tracks were color-coded based on LD r^2^ relative to the variant rs3740400. **(D)** Sashimi plots showing the gene model of *AS3MT* and the genomic position of unannotated splice site (blue) overlapping with *AS3MT* gene. **(E)** Average INT normalized PSI values for the splice site are shown for each genotype group. Schizophrenia risk allele T regulates the exon skipping event in progenitors. Boxplots showing INT normalized PSI values for splice across. **(F)** Genomic tracks color-coded based on pairwise LD r^2^ relative to the variant rs1222218 showing regional association of variants with SCZ and an unannotated alternative splicing event for *ARL14EP* gene in progenitors and neurons, association p-values on the y-axis, and genomic location of each variant on the x-axis. A cryptic exon skipping splice site (chr11:30323202-30332866) was associated with progenitor sSNP (rs1222218) colocalized with SCZ GWAS index SNP (rs1765142). **(G)** Sashimi plots with the gene model of *ARL14EP* and the genomic position of the unannotated splice site (blue) overlapping with *ARL14EP* gene. Average INT normalized PSI values for the splice site are shown for each genotype group. Schizophrenia risk allele G regulates the exon skipping event in progenitors. **(H)** Boxplots showing INT normalized PSI values for splice across rs1222218 genotypes in progenitors and neurons.

### Genetic imputation of cell-type specific GWAS susceptibility genes and alternative splicing

Next, we imputed genes and alternative splicing associated with brain related traits by integrating the polygenic impact of cell-type specific regulatory variants with GWAS risk variants in a transcriptome-wide association study (TWAS) approach ^52^. We found 1,703/973 genes and 6,728/6,799 intron junctions as significantly cis-heritable in progenitors/neurons (heritability *p-value* < 0.01). We found the cis-heritable impact of 124/102 genes and 359/365 intron junctions in progenitor/neuron significantly correlated with at least one brain related-traits (Table S5). Of those significant TWAS genes/introns, we separated conditionally independent genetic predictors from the co-expressed ones, and defined them as jointly independent ^53^. We performed cell-type specific TWAS on both gene expression and splicing for schizophrenia (jointly independent genes: 23/26; jointly independent introns: 59/64 in progenitor/neuron), IQ (jointly independent genes: 25/24; jointly independent introns: 38/65 in progenitor/neuron), and neuroticism (jointly independent genes: 13/15 neuron; jointly independent introns: 40/32 in progenitor/neuron) (Figure 5A-5C and Figure S5A-S5C). Also, we found novel loci not discovered in colocalization analysis per trait, demonstrating the additional power of TWAS compared to a single-marker testing approach. Despite the difference in population structure between our dataset and European neuropsychiatric GWAS, we observed that TWAS genes/introns were highly overlapped when different LD estimates were used (Figure S5E).

**Figure 5.**
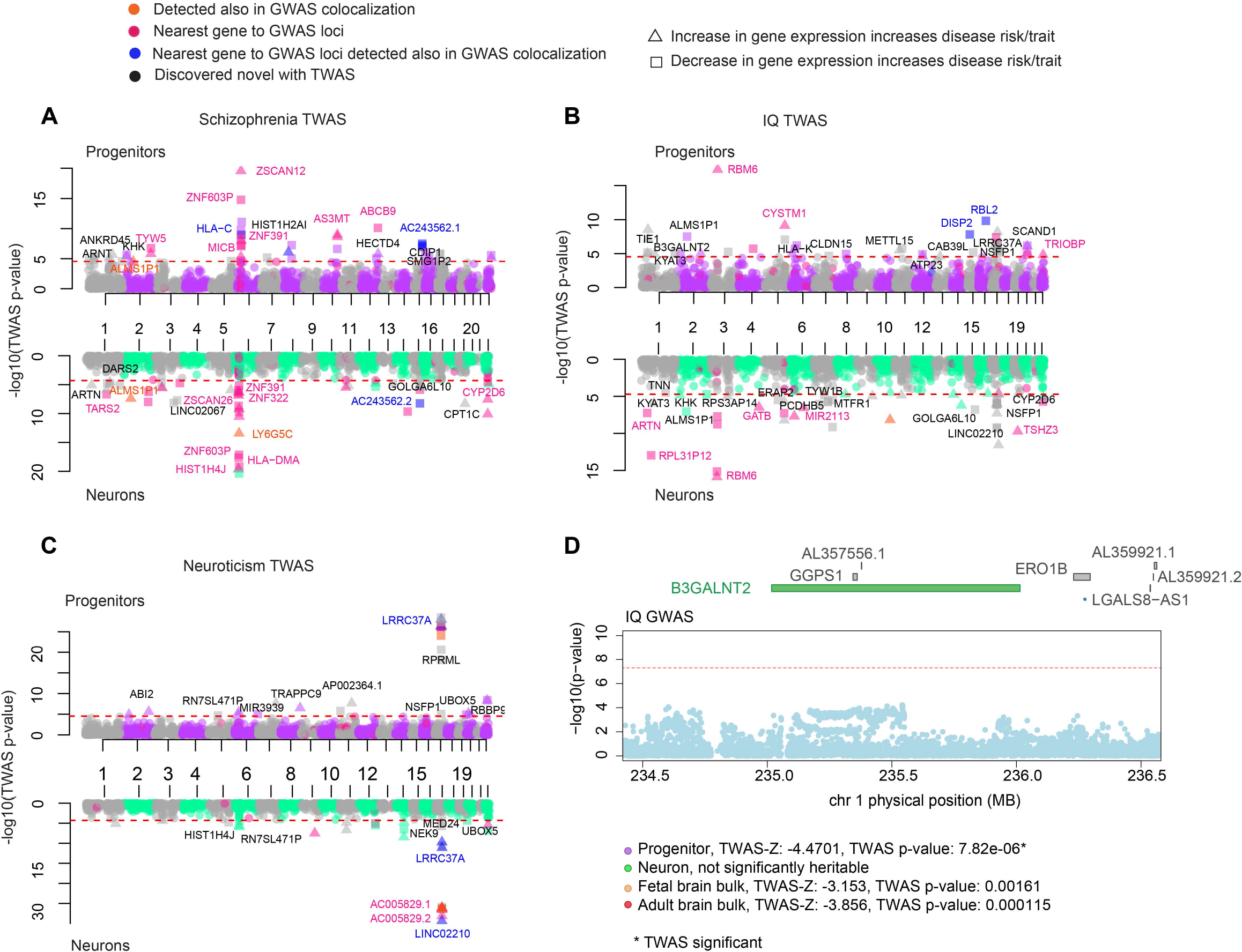
Prediction of gene expression during human brain development via TWAS. **(A)** Manhattan plots for schizophrenia, IQ, and neuroticism TWAS for progenitors (purple-grey, top) and neurons (green-grey, bottom) where the LD matrix used was based on a European population. Each dot shows −log10 (TWAS p-value) for each gene on the y-axis, genes were color-coded based on discovery also in colocalization analysis (orange), defined as the nearest gene to GWAS loci (dark pink), being in both these two categories (blue), and discovered only in TWAS analysis (black). Only joint independent genes are labelled (positively and negatively correlated genes represented by triangle and square, respectively and red line used for TWAS significant threshold) **(B)** Manhattan plots for IQ TWAS, as described in A. **(C)** Manhattan plots for neuroticism TWAS, as described in A. **(D)** IQ TWAS results for the *B3GALNT2* gene, regional association of variants to IQ trait shown at the top, and statistics from each TWAS study shown at the bottom (red line used for genome-wide significant threshold 5 x 10^-8^).

We next compared our cell-type specific TWAS approach to TWAS analyses performed using weights calculated from bulk cortical fetal tissue ^19^, and adult brain e/sQTLs from the Common Mind Consortium (CMC) ^14, 53^. Most TWAS findings were specific to a cell-type or temporal e/sQTL dataset, rather than broadly detected, indicating that different developmental or cell-type e/sQTL datasets contribute complementary information about genes influencing risk for neuropsychiatric disorders or other brain traits (Figure S5F and Table S5 for comparison). As an example, despite IQ GWAS falling short of the genome-wide significance threshold at *B3GALNT2* gene locus, we detected that genetically imputed *B3GALNT2* expression was significantly correlated with IQ in progenitors, but not in neuron, fetal bulk tissue or in CMC adult brain tissue (Figure 5D). Mutations in the *B3GALNT2* gene play a role in glycosylation of α-dystroglycan and were associated with intellectual disability in individuals with congenital muscular dystrophy ^54^. Overall, here we showed that an increase in *B3GALNT2* gene expression in progenitors is associated with lower IQ, suggesting this gene’s early cell-type specific impact on cognitive function.

Within the cell-type specific splicing TWAS, we found an intron junction of *MRM2* more frequently spliced that was associated with increased risk for schizophrenia specifically in progenitor cells (*TWAS-Z*: 6.538), but it was not significantly cis-heritable within neuron, fetal bulk or adult bulk data (Figure S5D). *MRM2* is a mitochondrial rRNA methyltransferase ^55^, and was found to be associated with intellectual disability ^56^ and mitochondrial encephalopathy ^55^. We propose a cell-type specific developmental basis for alternative splicing of the *MRM2* gene associated with risk for schizophrenia.

## Discussion

Here, we investigated the influence of genetic variation on brain related traits within a cell-type specific model system recapitulating a critical time period of human brain development, neurogenesis. Our analysis discovered features of gene regulation that will be complementary to previous eQTLs and sQTLs identified in bulk human brain in that: (1) we identified thousands of novel eQTLs, ASEs, and sQTLs during brain development that are enriched in regulatory elements present during neurogenesis; (2) most e/sQTLs in progenitors/neurons were not identified in previous fetal bulk post-mortem tissue datasets indicating the importance of cell-type specificity for identifying genetic influences on gene regulation; (3) using this resource, we are able to propose cell-type specific variant-gene/transcript-trait(s) pathways to further explore molecular and developmental causes of neuropsychiatric disorders; (4) by integrating the polygenic effects across traits and gene expression, we are able to impute cell type specific gene expression/alternative splicing dysregulation in individuals with neuropsychiatric disorders in time periods prior to disease onset.

As one example of a cell-type specific variant-gene-trait pathway, we discovered a locus near the *CENPW* gene colocalized across cell-type specific caQTL, eQTL, brain size, and cognitive function. Through the integration of multi-omic gene-to-trait databases, we hypothesize that the C allele at rs9388486 leads to decreased TF binding of up to 4 transcription factors (ATF1/2/4, CREM) in progenitors, resulting in decreased chromatin accessibility at the promoter peak, decreased expression of *CENPW*, leading to increased cortical surface area, and increased cognitive function. The *CENPW* gene has a strong role in proliferation, as it is required for kinetochore formation during mitosis ^57^. This is consistent with progenitor proliferation influencing surface area, as described in the radial unit hypothesis ^58^. Increased levels of *CENPW* may cause death of progenitor cells either by directly being an apoptotic inducer or by triggering apoptosis in response to an imbalance in cell homeostasis with excessive mitotic activity ^49^. In all, we demonstrate how integration across multi-level biological data can be used to propose functional mechanisms underlying complex traits, and future studies may be able to develop computational models to propose causal pathways across multi-omic QTL data ^9, 59, 60^. Such information will be crucial to both design efficient functional validation experiments as well as to leverage GWAS loci to advance treatment targets for neuropsychiatric disorders.

Though the most commonly proposed regulatory mechanism by which non-coding genetic variation influences complex traits is through gene expression levels ^32^, our data also support mechanisms by which genetic variants associated with cell-type specific alternative splicing influence complex brain-relevant traits. Importantly, we observed sQTLs impacting previously unannotated cell-type specific alternative splicing events that are also colocalized with brain relevant GWAS. For example, we found a progenitor specific sSNP regulating one unannotated exon skipping splice site for the *ARL14EP* gene also colocalized with an index SNP for schizophrenia GWAS, indicating a developmental molecular pathway contributing to schizophrenia risk.

Our cell-type specific TWAS analysis identified that alteration in expression of multiple genes and transcripts are associated with risk for different neuropsychiatric conditions. We followed a unique TWAS approach allowing us to explore cell-type and temporal specificity by leveraging existing fetal brain bulk and adult e/sQTLs together with the cell-type specific data we generated here. This type of analysis allows the imputation of the genetically regulated component of differential expression within cell types years prior to disease onset. As such, it allows the knowledge of gene expression differences that cannot be gained from post-mortem tissue of cases versus controls, which must be acquired after diagnosis. This window into developmental gene expression differences may be particularly important to understand disease risk, as these results are not subject to confounding by medication use or the altered experiences of the environment of individuals living with a neuropsychiatric illness ^61^. Nevertheless, further support for such data could be gained from iPSC lines modeling early developmental time periods from large populations of cases vs controls.

With our cell-type specific model, we propose how and when genetics influence brain related traits through gene expression and splicing. Future cell-type specific e/sQTLs acquired using flow cytometry or scRNA-seq from the developing post-mortem brain will be useful to validate the *in vivo* impact of genetic variants discovered here using an *in vitro* system. Nevertheless, this *in vitro* system has particular utility in that, in the future it may be used to determine the impact of genetic variation in response to activation of specific pathways or response to environmental stimuli ^23^. By pursuing cell-type, temporal, and environmental specificity of e/sQTLs, we expect that a greater degree of mechanisms underlying risk for neuropsychiatric disorders and brain-relevant traits can be uncovered.

## Acknowledgments

This work was supported by NIH (R00MH102357, U54EB020403, R01MH118349, R01MH120125), Brain Research Foundation, and NC TraCS Pilot funding to JLS. DHG was supported by NIH (R37 MH060233, R01 MH094714, UO1MH116489 and R01 MH110927). The following core facilities were utilized for this project: UNC Neuroscience Center Microscopy Core (P30NS045892), UNC Mammalian Genotyping Core, CGIBD Advanced Analytics Core (NIH grant P30 DK034987), UNC Flow Cytometry Core Facility, UNC Vector Core, UNC Research Computing. Additional core facilities utilized for this project were: UCLA CFAR (5P30 AI028697), and the UCLA Neuroscience Genomics Core. We thank Dr. Karen L. Mohlke and Dr. Yun Li for helpful comments, Dr. Eric Wexler for the idea of phNPC eQTL, and Dr. Stephen Montgomery for clarifying the eigenMT method.

## Author Contributions

JLS, DHG, and LTU conceived the study. JLS directed and supervised the study. JLS along with DHG provided funding. ALE, KEC, KPC, MY, LTU, and JLS cultured human neural progenitor cells. ALE performed library preparation. MJL and NA pre-processed the RNA-seq data. DL performed pre-processing of genotyping data and caQTL analysis. JM, EFH and OK performed immunocytochemistry. MIL aided in differential gene expression and allele specific expression analysis methodology. NA managed the integration of the datasets, interpreted results and performed TMAP, differential gene expression and splicing, eQTL, ASE, sQTL, co-localization, enrichment, transcription factor binding motif, and TWAS analyses. JLS and NA wrote the manuscript. All authors commented on and approved the final version of the manuscript.

## Data and code availability

Data will be available within to dbGaP upon publication, and codes are available here https://bitbucket.org/steinlabunc/expression_splicing_qtls_public/src/master/.

## Supplemental Information

**Figure S1, related to Figure 1:**
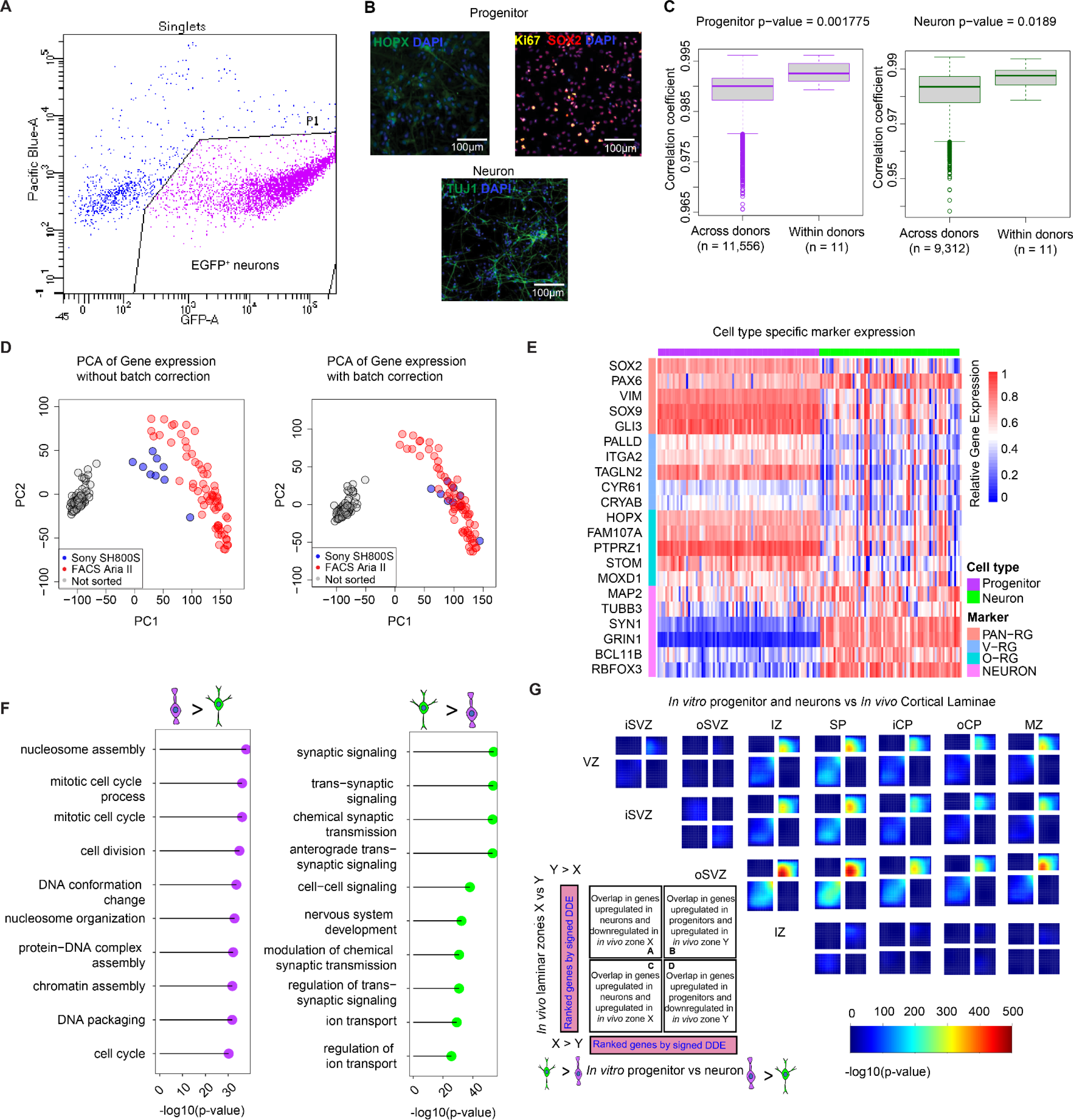
Pre-processing RNA-seq data and evaluation of the fidelity of *in vitro* cell-type specific system. **(A)** Flow cytometry results showing sorting of live EGFP positive neurons in pink. The y-axis marks fluorescence from a live/dead stain (annexin V/SYTOX) and the x-axis marks fluorescence from GFP. **(B)** Immunolabeling indicates outer radial glia marker HOPX in green, proliferation marker Ki67 in yellow and pan-radial glia marker SOX2 in red were expressed in undifferentiated progenitor cultures, and the neuronal marker TUJ1 was expressed in neurons from 8 week differentiated cultures (scale bar is 100 μm, DAPI in blue). **(C)** Replicate correlation of RNA-seq libraries across donors and within donors. Gene expression profiles were more correlated between libraries generated from the same donor thawed at different times as compared to libraries across different donors for both progenitors (left, p-value=0.001775) and neurons (right, p-value=0.0189). **(D)** Principal component analysis (PCA) before and after batch correction of neuron for the machine (Sony SH800S in blue, FACS Aria II in red, progenitors not sorted in grey) used for sorting. **(E)** Heatmap showing cell-type specific expression of literature based progenitor (PAN-RG: Pan-radial glia, V-RG: ventricular radial glia, O-RG: outer radial glia) and neuronal markers listed on the y-axis. The x-axis indicates progenitor (purple) or neuron (green) cells from each donor. The color of the heatmap indicates the relative gene expression normalized for each gene between 0 and 1. **(F)** Gene ontology (GO) analysis showing pathways enriched for genes upregulated in progenitors (left, in purple), and for genes upregulated in neurons (right, in green). The x-axis shows adjusted −log10(p-values) for enrichment and each GO term is listed in the y-axis. **(G)** Comparison of the transitions between mitotic and postmitotic regions of *in vivo* cortical laminae in the developing cortex and in vitro progenitor and neurons with rank-rank hypergeometric overlap (RRHO) maps. The extent of overlap between *in vivo* and *in vitro* transcriptome was represented by each heatmap colored based on −log10(p) value from a hypergeometric test. Each map shows the extent of overlapped upregulated genes in the bottom left corner, whereas shared downregulated genes are displayed in the top right corners (ventricular zone - VZ; inner and outer subventricular zone - i/oSVZ, intermediate zone - IZ; subplate - SP; inner and outer cortical plate - i/oCP, marginal zone - MZ).

**Figure S2, related to Figure 1, Figure 2 and Methods:**
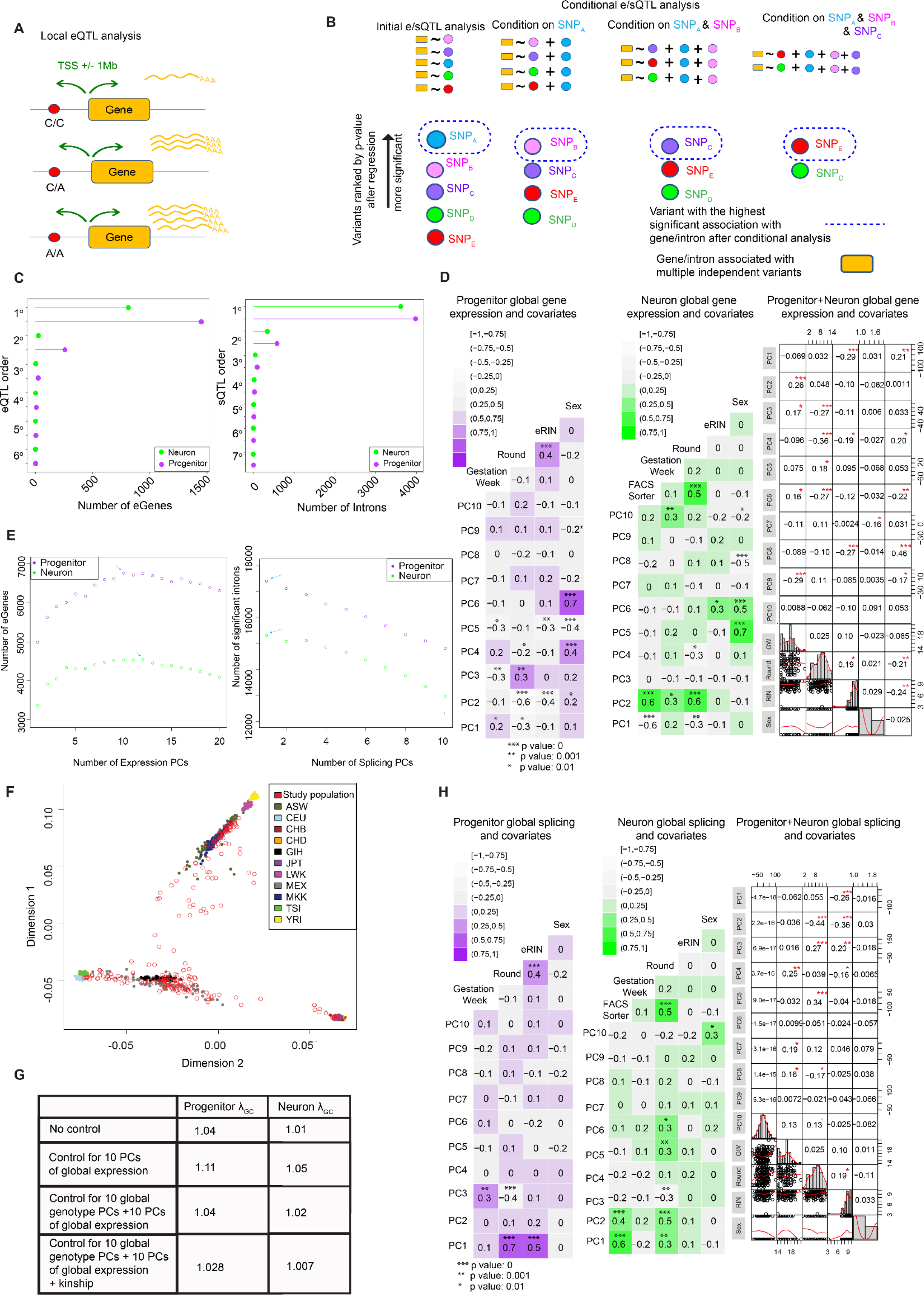
Local eQTL/conditional analysis design and the detection of covariates for e/sQTLs. **(A)** A schematic showing that variants within +/− 1MB cis window from the transcription start site (TSS) of each gene were tested for the association with gene expression. **(B)** A schematic showing the conditional e/sQTL procedure. Conditionally independent SNPs were found conditioning on the genetic variant with the most significant association, and iteratively applying the same algorithm until there were no further significant associations with local variants. **(C)** Number of eGenes on the x-axis regulated by number of conditionally independent eSNPs on the y-axis indicated by eQTL order (left). Number of intron junctions on the x-axis regulated by number of conditionally independent sSNPs on the y-axis indicated by sQTL order (right). **(D)** Correlation of technical confounders with the top 10 principal components of gene expression in progenitor, neurons and all data (asterisk indicates significant correlation). **(E)** Covariate selection analysis for eQTLs with number of eGene vs. number of global gene expression PCs (left, progenitors in purple, neurons in green). Covariate selection analysis for sQTLs with number of significant intron vs. number of global splicing PCs (right, progenitors in purple, neurons in green). Blue arrows indicate the number of PCs used in each dataset. **(F)** Multidimensional scaling (MDS) of global genotypes showing the multi-ancestry donors in our study. MDS1 vs MDS2 values plotted where each red circle represents a unique donor in our study and each different color represents different ancestry from HapMap3 (ASW: African ancestry, CEU:Northern and Western European ancestry, CHB: Han Chinese ancestry, CHD: Chinese in metropolitan Denver, GIH: Gujarati Indians in Houston, JPT: Japanese in Tokyo, LWK: Luhya in Webuye, MEX:Mexican ancestry, MKK: Maasai in Kinyawa, TSI: Toscani in Italy, YRI: Yoruba in Ibadan). **(G)** Comparison of genomic inflation factor (λ_GC_) without controlling for population structure and technical confounders (no control), only controlling for technical confounders by adding global gene expression PCs, controlling for both population structure (10 MDS of global genotype) and technical confounders, and controlling for kinship matrix in addition to the previous covariates. **(H)** Correlation of technical confounders with top 10 principal components of splicing in progenitor, neurons and all data (asterisk indicates significant correlation).

**Figure S3, related to Figure 1:**
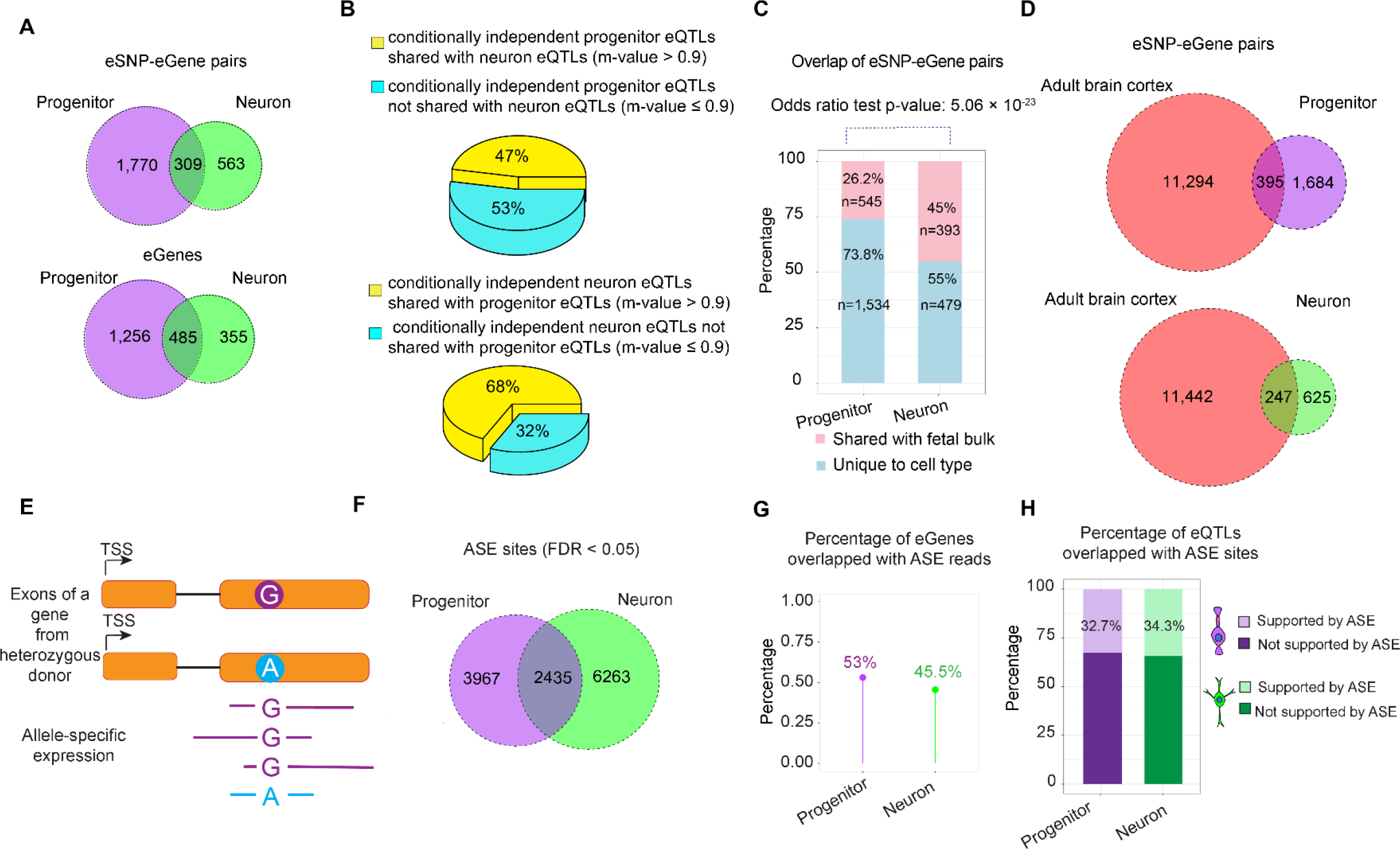
Cell-type and temporal specificity of eQTLs and ASE analysis. **(A)** Overlap between progenitor and neuron eQTLs for eSNP-eGene pairs and eGenes. **(B)** Posterior probability of shared effect size (m-value). Upper pie shows the percentage of conditionally independent progenitor eQTLs shared with neuron eQTLs, and lower pie shows the percentage of conditionally independent neuron eQTLs shared with progenitor eQTLs at m-value > 0.9. **(C)** Overlap percentage of cell-type specific eSNP-eGene pairs shared (pink) with fetal bulk eQTLs (variants with LD r^2^ > 0.8 were considered as the same loci). Odds ratio test p-value is shown. **(D)** Overlap between progenitor/neuron eQTLs and adult brain cortex eQTLs for eSNP-eGene pairs. **(E)** A schematic illustrating allele specific expression (ASE) in a heterozygous individual for a variant of interest. **(F)** Overlap between progenitor and neuron specific ASE sites. **(G)** Overlap between eGenes and genes with ASE (progenitors in purple, neurons in green). **(H)** Overlap between cell-type specific eSNPs and ASE sites (progenitors in purple, neurons in green).

**Figure S4, related to Figure 2:**
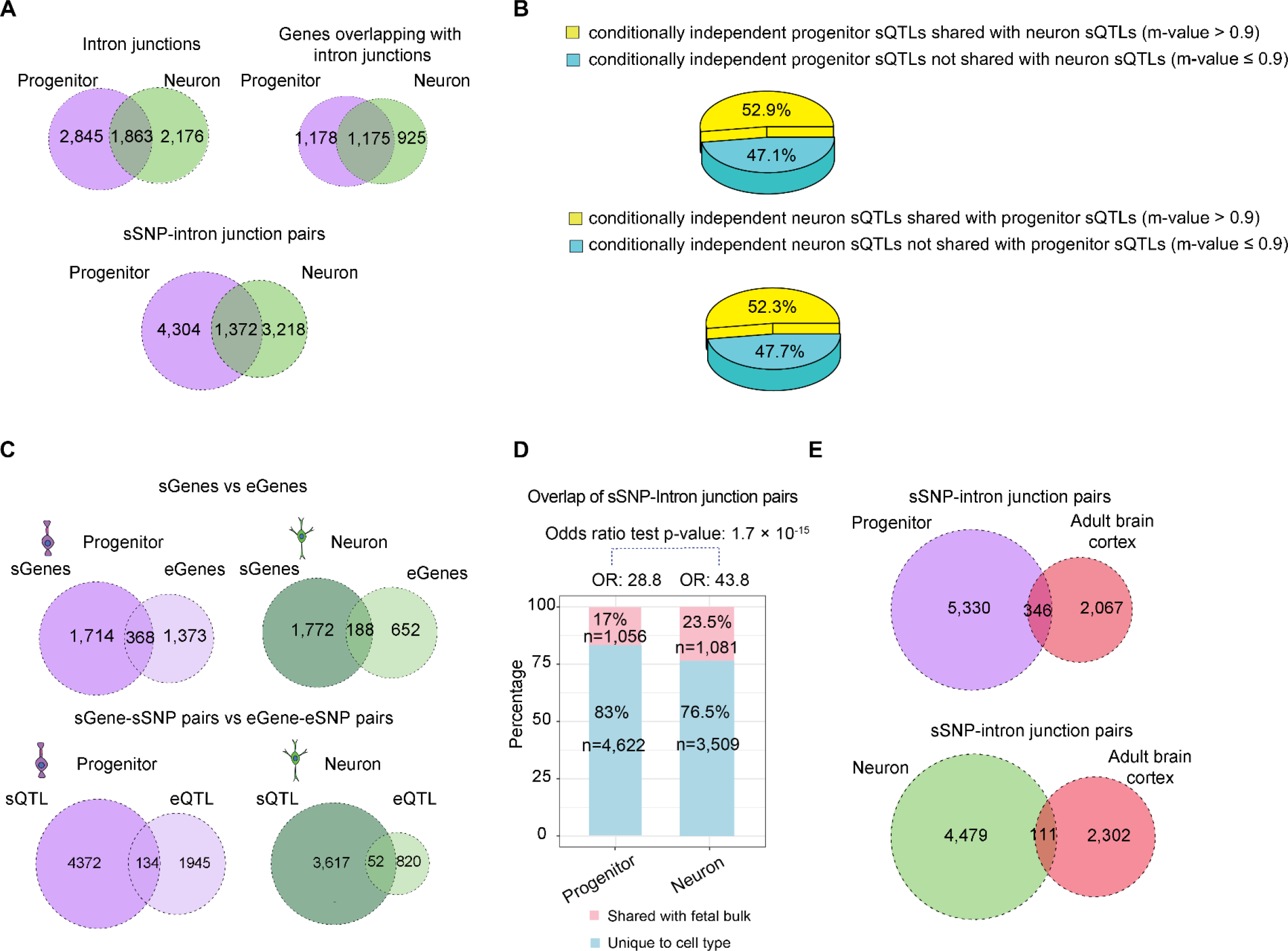
Cell-type and temporal specificity of sQTLs. **(A)** Overlap of intron junctions, sGenes that are the genes intron junctions span and sSNP-intron junction pairs for progenitor vs neuron sQTLs **(B)** Posterior probability of shared effect size (m-value). Upper pie shows the percentage of conditionally independent progenitor sQTLs shared with neuron sQTLs, and lower pie shows the percentage of conditionally independent neuron sQTLs shared with progenitor sQTLs at m-value > 0.9. **(C)** Comparison of cell-type specific sQTL vs eQTLs, progenitor in purple and neuron in green. Overlap between sGenes and eGenes, upper panel; overlap between sGene-sSNP and eGene-eSNP pairs, lower panel. **(D)** Overlap percentage of cell-type specific sSNP-intron junction pairs shared (pink) with fetal bulk sQTLs (variants with LD r^2^ > 0.8 were considered as the same loci). Odds ratio test p-value is shown **(E)** Overlap between progenitor (in purple)/neuron (in green) sQTLs and adult brain cortex sQTLs (in red) for intron junction-sSNP pairs.

**Figure S5, related to Figure 5:**
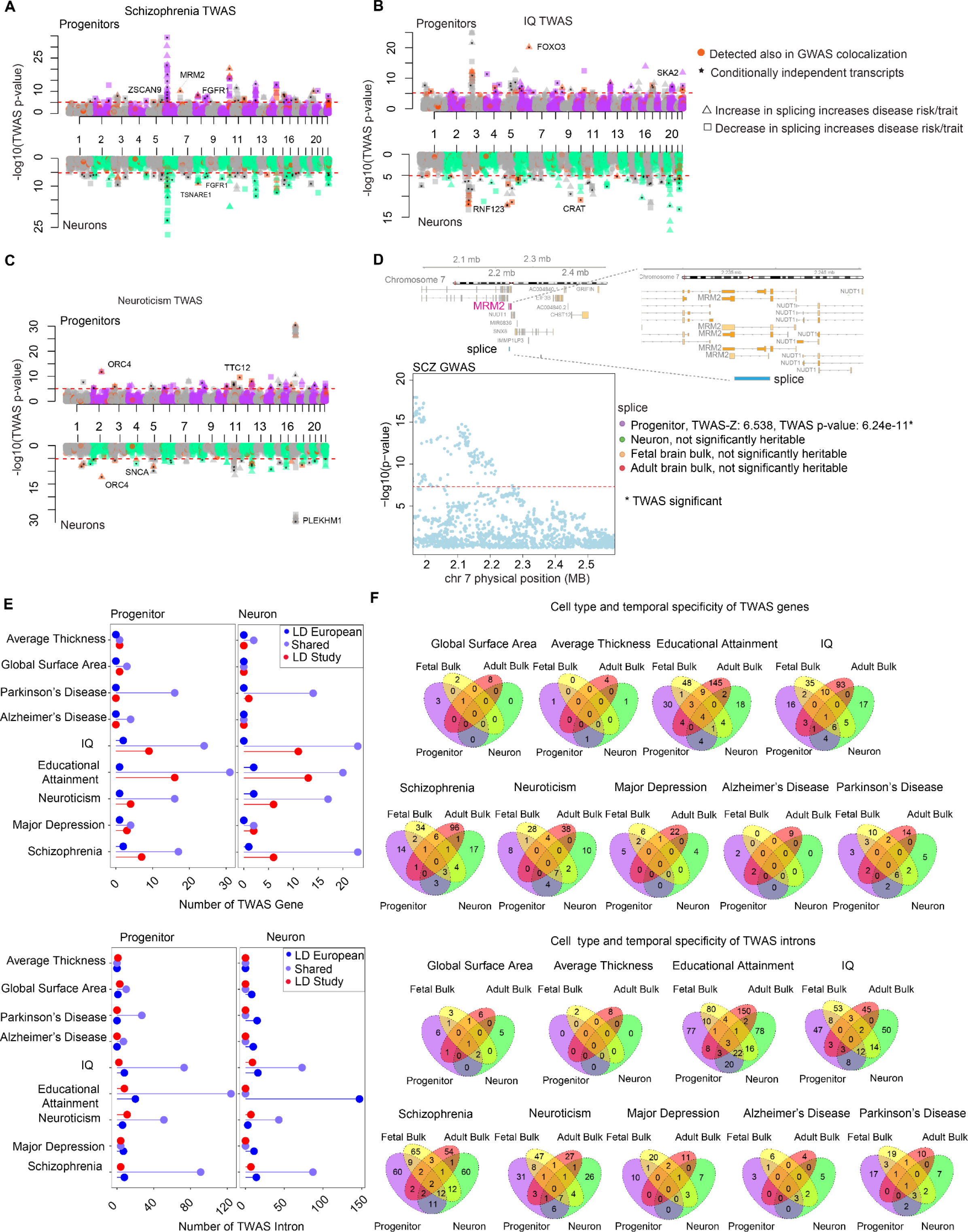
Prediction of alternative splicing events during human brain development via TWAS, and cell-type/temporal specificity of TWAS genes and introns. **(A)** Manhattan plots for schizophrenia TWAS for progenitors (purple-grey, top) and neurons (green-grey, bottom) where LD matrix calculated based on a European population. Each dot shows the −log10(TWAS p-value) for each intron junctions on the y-axis, introns were color-coded based on discovery also in colocalization analysis (orange), and being jointly independent (asteriks), where positively and negatively correlated splicing represented by triangle and square, respectively. **(B)** Manhattan plots for IQ TWAS with graphic design described in A. **(C)** Manhattan plots for Neuroticism TWAS with graphic design described in A. **(D)** SCZ TWAS results for intron junction (splice, chr7:2235564-2239418) of the *MRM2* gene, regional association of variants, that were used to test polygenic impact on introns to SCZ are shown on the left. Gene-model for *MRM2* is shown on the right with matching introns and statistics from each TWAS study shown at the bottom (red line used for genome-wide significant threshold of 5 x 10^-8^). **(E)** Comparison of TWAS genes performed by using different LD matrices based on European (LD European) and population included in our QTL study (LD Study) (upper plot). Comparison of TWAS introns performed by using different LD matrices based on European (LD European) and population included in our QTL study (LD Study) (lower plot). **(F)** Overlap of cell-type specific TWAS genes (from the analysis where LD was estimated from European population) with fetal brain bulk and adult brain bulk TWAS genes (upper plot). Overlap of cell-type specific TWAS introns (from the analysis where LD was estimated from European population) with fetal brain bulk and adult brain bulk TWAS introns (lower plot).

## Supplementary Table legends

**Table S1**, related to Figure 1, S1, S2 and S3:

Sheet 1: Differential gene expression analysis progenitor vs neurons (FDR < 0.05): gene is the ensemblID, logFC is the expression fold change logFC > 0 indicates a gene more frequently expressed in neurons than progenitors; AveExpr is the average vst normalized expression of all samples. t is the expression fold change divided by its standard error ^35^. P.Value is the nominal p-value from the testing differential expression; adj.P.Val is the Benjamini-Hochberg FDR adjusted p-value; B is log-odds for the differentially expressed gene in limma.

Sheet 2-4: List of cell-type specific conditionally independent eQTLs for progenitor, neurons and fetal bulk: snp is the variant tested in QTL; beta is the beta coefficient; pval is the nominal p-value; gene is the ensemblID of the gene tested; rank is the eQTL order; chr is the chromosome number, BP is the genomic position of the variant; Cond.beta is the beta after conditional analysis; Cond.pval is the p-value after conditional analysis; A1 is the effect allele.

**Table S2,** related to Figure 1 and S3F: Allele specific expression analysis (FDR < 0.05). SNP is the variant tested for allele specific expression analysis, baseMean is the average of the normalized count values divided by size factors from DESeq2 ^71^; log2FoldChange is the expression fold change logFC > 0 indicates reads more frequently expressed in donors with reference allele than donors with alternative allele; lfcSE is the standard error estimate for log2FoldChange; stat is the test statistics performed in DESEq2; pvalue is the nominal p-value from the testing differential expression; padj is the Benjamini-Hochberg FDR adjusted p-value; refAllele is the reference allele of the variant.

**Table S3,** related to Figure 2 and S4:

Sheet 1: Differential splicing analysis progenitor vs neurons (FDR < 0.05): intron is the splice junction, logFC is the expression fold change logFC > 0 indicates a gene more frequently expressed in neurons than progenitors; AveExpr is the average vst normalized expression of all samples. t is the expression fold change divided by its standard error ^35^. P.Value is the nominal p-value from the testing differential expression; adj.P.Val is the Benjamini-Hochberg FDR adjusted p-value; B is log-odds for the differentially expressed intron in limma; chr is the chromosome, start is the start position of the junction; end is the end position of the junction; clusterID is the cluster identified from Leafcutter, cluster is the clusterID combined with chromosome number, verdict is the annotation status; gene is the gene symbol of the gene that introns junctions overlap with; ensemblID is the ensemblID of that gene; transcripts is the transcripts where intron junction overlap with; constitutive.score: degree of the junction shown in each transcript.

Sheet 2-4: List of cell-type specific conditionally independent sQTLs for progenitor, neuron and fetal bulk sQTLs: snp is the variant tested; beta is the beta coefficient, pval is the nominal p-value; intron is the intron junction as chromosome:start position:end position format; rank is the order of sQTL after conditional analysis; chr is the chromosome, start is the start position of the junction; end is the end position of the junction; clusterID is the cluster identified from Leafcutter, cluster is the clusterID combined with chromosome number, verdict is the annotation status; gene is the gene symbol of the gene that introns junctions overlap with; ensemblID is the ensemblID of that gene; transcripts is the transcripts where intron junction overlap with; constitutive.score: degree of the junction shown in each transcript; Cond.beta is the beta coefficient after conditional analysis (for primary QTLs, it is identical to beta); Cond.pval is the p-value after conditional analysis (for primary QTLs, it is identical to pval), A1 is the effect allele; rsid is the rs id of the allele matching in 1000 Genome Phase 3.

Sheet 5: Enrichment of RNA binding protein (RBP) sites within cell-type specific sQTLs. PThresh is the p-value threshold used for enrichment; OR is the odd ratio; Pvalue is enrichment p-value; Beta is the beta coefficient after enrichment test via GARFIELD ^83^; SE is the standard error; CI95_lower is the lower bound of 95% confidence interval; CI95_upper is the upper bound of 95% confidence interval; NAnnotThesh is the is the number of annotated variants at the p-value threshold; NAnnot is the total number of variants after pruning; NThresh is the number of variant passing p-value threshold after pruning; N is the number of variants remained after pruning; linkID is the ID in annotation file; Annotation is the RNA-binding protein; Tissue is the the type of tissue used; Type is the cell type used for enrichment test.

**Table S4,** related to Figure 3 and 4: Colocalization of GWAS for neuropsychiatric disease and other brain related traits with cell-type specific e/sQTLs and fetal bulk e/sQTLs: e/sQTLsnp is the e/sSNP; inibeta is the beta coefficient before conditioning on GWAS SNP; pval is the nominal p-value prior to conditional analysis, gene/intron is the ensemblID of gene/intron junction associated with the e/sSNP; A1 is the effect allele for e/sQTL index SNP; GWASsnp is the variant e/sSNP colocalized with; Condbeta is the beta estimate of e/sQTL after conditional analysis; Condpval is the p-value after conditional analysis; r2 is the linkage disequilibrium (LD) r^2^; pop is the population used to estimate LD r^2^ (European population, with “European” or the population used in the QTL study with “Study”); symbol of the symbol of the gene (for eQTLs); biotype is the biotype of the gene for eQTLs; trait is the trait for GWAS.

**Table S5,** related to Figure 5 and S5:

Sheet 1-8: List of cell-type specific/fetal bulk/adult bulk TWAS gene and introns for neuropsychiatric disease and other brain related traits. Output from FUSION ^52^ : ID is the gene ensemblID or intron id; CHR is the chromosome number; HSQ is the heritability; BEST.GWAS.ID is the GWAS SNP in the locus with the most significant association; BEST.GWAS.Z is the z-score of the best GWAS SNP; EQTL.ID is the best e/sQTL in the locus; EQTL.R2 is the cross-validation R^2^ of the best e/sQTL in the locus; EQTL.Z is the z-score of the best e/sQTL in the locus; EQTL.GWAS.Z is the GWAS Z-score for this e/sQTL; NSNP is the number of SNPs in the locus; NWGT is the number of snps with non-zero weights; MODEL is the best performing model; MODELCV.R2 is the the cross-validation R^2^ of the best performing model; MODELCV.PV is the p-value from the cross-validation of the best performing model; TWAS.Z is the TWAS z-score; TWAS.P is the TWAS p-value; trait is the GWAS trait; pop is the population used to estimate LD; joint_independent is the status if a gene/intron jointly independent (YES, if it is independent; NO, if it is not independent; NA, if it was not tested for the trait).

Sheet 9-10: Summary of heritability (p-value < 0.01) and cross validation r^2^ from prediction models across cell-type specific/fetal bulk/adult bulk for gene and intron TWAS: hsq is the mean heritability of the genes/introns; hsq.se is the mean standard error of estimated heritability; hsq.pv is the mean p-value of the heritability; emmax.rsq is the mean cross-validation R^2^ training via EMMAX with p-value as emmax.pval; lasso.rsq is mean the cross-validation R^2^ via LASSO with p-value as lasso.pval; enet.rsq is mean the cross-validation R^2^ via elastic net with p-value as enet.pval; blup.rsq is mean the cross-validation R^2^ via BLUP with p-value as blup.pval; bslmm.rsq is the mean cross-validation R^2^ via BSLMM with p-value as bslmm.pval; top1.rsq is the mean cross-validation R^2^ via standard marginal e/sQTL Z-scores computation with p-value as top1.pval. 95 % confidence intervals per parameter are shown their below.

## Material and Methods

### Cell Culture

Generation of human neural progenitor cells was previously described^16, 26^. Briefly, human fetal brain tissue was acquired from the UCLA Gene and Cell Therapy Core following IRB regulations from approximately 14-21 gestation weeks (inferred to be 12-19 postconceptional weeks). Presumably cortical tissue was selected by visual inspection, subjected to single cell dissociation, and cultured as neurospheres. Neurospheres were plated on laminin/fibronectin and polyornithine coated plates for an average of 2.5 ± 1.8 s.d. passages, and cryopreserved as primary human neural progenitors (phNPCs).

Cryopreserved phNPCs were transferred to UNC Chapel Hill, after material transfer agreement, where all downstream culture and analyses were completed. Donors processed for ATAC-seq (described previously ^16^) and RNA-seq (described here) were cultured simultaneously. The overall design of the experiment and media used for culture was previously described ^16^. Briefly, we cultured 89 unique donors for subsequent RNA-seq library preparation. We first randomly assigned the approximately 8-9 donors into 10-12 rounds for a feasible cell culture workload. We thawed one round every three weeks. To reduce batch effects, we processed each round on the same day of the week and designated the same person to do each task as much as possible. Cells were isolated at two time points: progenitor and their differentiated and virally labeled neuronal progeny. Progenitors were cultured in proliferation media including growth factors for 3 weeks, and we lifted them with trypsin to prepare RNA-seq libraries. Differentiation was performed for 5 weeks, after which the culture was transduced with AAV2-hSyn1-eGFP (https://www.addgene.org/50465/) virus at 20,000 multiplicity of infection (MOI) to label neurons and then differentiated for another 3 weeks. FACS sorting (using either BD FACS Aria II or Sony SH800S) at 84 days post-differentiation was used to isolate EGFP labelled neurons (Figure S1A). After cells were isolated as either progenitors or neurons, we added Qiazol and stored the mixture at −80°C for randomized RNA isolation to reduce batch effects.

### Immunofluorescence labeling and imaging

At the progenitor stage or after 8 weeks of differentiation, we fixed the cells by incubating them in 4% PFA, and performed permeabilization with 0.4% Triton in PBST. We used 10% goat serum dissolved in PBST for blocking. We incubated blocked samples with primary antibodies dissolved in PBST solution with 3% goat serum at 4°C overnight followed by washing 3 times with PBST. Samples were subject to incubation in fluorophore-conjugated secondary antibodies, for 1 hour at room temperature, then they were stained with DNA-binding dye DAPI with 10 minutes incubation. We used antibodies with concentrations listed as following: SOX2 (1:400, rabbit, Millipore #AB5603), Ki67 (1:1000, rat, Invitrogen #14-5698-82), TUJ1 (1:2000, mouse, Biolegend #801202), Alexa Fluor 568 (1:1000, goat anti-rabbit, Invitrogen #A11036), Alexa Fluor 647 (1:1000, goat anti-rat, Invitrogen #A21247), Alexa Fluor 488 (1:1000, goat anti-mouse, Invitrogen #A11001).

### RNA-seq Library Preparation

We isolated RNA from progenitors and neurons using Qiagen miRNeasy Minelute kit, quantified RNA concentration with a Qubit 2.0 fluorometer, and assessed RNA integrity via eRIN scores using the Agilent Tapestation. We prepared libraries for sequencing using Kapa Biosystems KAPA Stranded RNA-seq with Riboerase (HMR) kit by loading 50 ng of total RNA into the initial reaction. We followed the manufacturer’s instructions for fragmentation and PCR steps. To obtain ∼350 bp average insert size, we fragmented cDNA at 85°C for 6 min. Final library concentrations were determined using Qubit 2.0 fluorometer and pooled to a normalized input library. Pools were sequenced on a NovaSeq S2 flowcell using 150 bp PE reads with an average read depth of 99M per sample.

### RNA-sequencing data processing

We merged fastq files from the same library when sequenced on multiple flow cells and trimmed the adapters using sequences provided by Illumina with Cutadapt/1.15^62^. Quality control of each library was performed with FastQC (https://www.bioinformatics.babraham.ac.uk/projects/fastqc/). For alignment, we first integrated the sequence of AAV2-hSyn1-eGFP plasmid (https://www.addgene.org/50465/) used for labeling neurons into GRCh38 release92 reference genome (https://www.ncbi.nlm.nih.gov/assembly/GCF_000001405.38/). Then, we aligned the fastq files to this combined reference genome by implementing STAR/2.6.0a aligner ^63^.

We processed aligned data further with different steps based on downstream analyses. To estimate gene expression levels, we quantified reads with the union exon based approach using featureCounts, where for each gene, all overlapping exons were merged to form union exons, and the reads mapped to those union exons with the same strandedness were counted ^64^. Gene models were identified using the GTF file Homo_sapiens.GRCh38.92 (http://ftp.ensembl.org/pub/release-92/gtf/homo_sapiens/Homo_sapiens.GRCh38.92.gtf.gz) merged with AAV2-hSyn1-eGFP plasmid.

For allele specific expression and splicing quantification, we remapped the aligned data with WASP software(v2018-07)^65^ to reduce reference mapping bias. First, we identified reads overlapping with bi-allelic SNPs within our acquired genotype data. Following this, the genotype of any reads overlapping with a SNP was swapped with the other allele, and re-mapped. WASP discarded re-mapped reads that did not map to the same genomic position. As a final step, we implemented *rmdup.py* script provided in the WASP software which removes duplicate reads randomly, regardless of their mapping score.

### Mycoplasma contamination test

Adapter trimmed reads (see above) were mapped using STAR to a combined reference including the GRCh38 release 92 human reference genome, AAV2-hSyn1-eGFP plasmid, and over 1400 mycoplasma genomes. Alignment parameters allowed for simultaneous mapping of reads to one or more human and mycoplasma genomes. No sample exceeded 0.11% of total reads mapping to any mycoplasma genome, indicating none of our cultures were contaminated with mycoplasma. This mapping was only used for mycoplasma contamination analysis and not for subsequent analyses.

### Genotype processing

We performed genotyping using Illumina HumanOmni2.5 or HumanOmni2.5Exome platform, and exported SNP genotypes to PLINK format following the procedure previously described ^16^. Briefly, we converted SNP marker names from Illumina KGP IDs to rsIDs using the conversion file provided by Illumina. We performed quality control with PLINK v1.90b3 software ^66^ as follows. We filtered out SNPs with the following criteria: variant missing genotype rate > 5% (--geno 0.05), deviations from Hardy-Weinberg equilibrium at p<1×10^-6^ (--hwe 10^-6^), minor allele frequency > 1% (--maf 0.01). We also filtered out individuals with missing genotype rate > 10% (--mind 0.10). We obtained 1,760,704 directly genotyped variants surviving our QC procedure. Lastly, we called sex from genotype data using PLINK v1.90b3 software based on heterozygosity on the X chromosome. When there was an ambiguity for sex assessment based on genotype data, we checked Xist gene expression. We estimated the population structure of our study cohort by implementing multidimensional scaling (MDS) for genotype data of our samples and genotype data from HapMap3 (https://www.sanger.ac.uk/resources/downloads/human/hapmap3.html). We followed the protocol from ENIGMA consortium (http://enigma.ini.usc.edu/wp-content/uploads/2012/07/ENIGMA2_1KGP_cookbook_v3.pdf). By plotting MDS1 vs MDS2, we visually show each donor’s ancestry relative to known populations (Figure S2F).

### Imputation

After filtering genotype data, we pre-phased the data with SHAPEIT v2.837 ^67^. For our imputation reference panel, we used 1000 Genomes Project Phase 3 that contains a total of 37.9 million SNPs in 2,504 individuals with multiple ancestries, including those from West Africa, East Asia and Europe ^68^). Imputation was implemented using Minimac4 software ^69^ (v1.0.0). On the X chromosome, we separately performed pre-phasing and imputation steps for the pseudoautosomal region and non-pseudoautosomal regions. Following imputation, we retained any variants with missing genotype rate lower than 0.05, Hardy-Weinberg equilibrium p-value lower than 1 x 10^-6^ and minor allele frequency (MAF) bigger than 1%. We retained SNPs with sufficient imputation quality (R^2^ > 0.3), and obtained approximately 13.6 million SNPs in total.

### Sample quality control

One library with missing eRIN score and one library with missing final cDNA concentration from neurons were removed. In order to detect sample swaps or mixing between samples, we evaluated consistency of genotypes called from the RNA-seq and genotyping array via VerifyBamID v1.1.3 ^70^. We removed RNA-seq libraries file with [FREEMIX] > 0.04 or [CHIPMIX] > 0.04 (N_library_ = 8). Also, we corrected samples where we detected swaps (N_library_= 7). After quality control, we retained 85 unique donors for progenitors, and 74 unique donors for neurons for subsequent analyses.

### Replicate correlation and determination of technical factors correlating with gene expression

Quantified RNA-seq reads with featureCounts were imported to generate a gene count matrix in DESeqDataSet format from DESeq2 R package ^71^. We filtered out the lowly expressed genes (those where fewer than 10 read counts of a gene were observed in fewer than 5% of samples), and normalized the data via variance stabilizing transformation (vst()) function from DESeq2 R package ^71^. We included genes on the X and Y chromosomes and genes transcribed from mitochondrial DNA meeting the expression criteria. We subset the normalized gene expression matrix into progenitor and neuron specific samples. To identify major axes of variation in gene expression across samples, we computed principal components of gene expression with prcomp() function from stats R package for each cell-type separately, and reported the proportion of variance explained by each component.

We recorded biological and technical variables for each sample which may potentially impact gene expression: cell type, postconception week, sex, tissue acquisition date, researcher extracting RNA and preparing libraries, RNA input amount, index number and bases, final cDNA concentration, BioAnalyzer run date, average basepair of BioAnalyzer cDNA, sequencing pool, cell input, Qiazol lot number and addition date, eRIN, RNA extraction date, RNA tapestation date, Qiagen extraction kit lot number, FACS sorting date and time, total live cells during sorting, FACS machines used, researcher performing FACS sorting, papain lot number and addition date, differentiation rank (a qualitative assessment of cell health evaluated under the microscope), well location in the 6-well plate, date to plate for differentiation, researcher washing and differentiating cells and date, virus addition date, researcher adding virus, PBS lot number used for cell proliferation and differentiation, laminin, polyornithine lot numbers used for proliferation and differentiation, donor ID, round, media lot numbers used for proliferation, passage number, split dates, researcher performing each split, rank for proliferation (qualitative assessment of cell health), trypsin lot number used for splitting cells, and fibronectin lot number. To identify technical covariates impacting expression levels, we assessed if any recorded biological or technical variables were significantly correlating with first 10 expression PCs separately for each cell type. We observed that different FACS machines (Sony SH800S with N_donor_ = 8; FACS Aria II with N_donor_= 66) used to isolate GFP labelled neurons had a strong impact on global gene expression in neurons (PC1: r = 0.59, p-value = 1.782e-08; PC2: r = 0.58, p-value = 3.972e-08) (Figure S2D). To remove the impact of sorter on global neuron expression profiles prior to differential expression analysis, we implemented function ^72^. Then, we combined the gene expression matrix from batch corrected neurons with progenitors gene expression data.

We cultured several donors multiple times during the course of the experiment in order to quantify cell culture induced noise. We calculated Pearson’s correlation of gene expression between libraries from the same donors (N_correlation_ = 11 in both progenitors and neurons), and between each library across donors in a pairwise manner (N_correlation_ = 11,556 for progenitors; N_correlation_ = 9,312 for neurons). For neurons, we used gene expression values after batch correction with the limma R package for the sorter type, as described above. We performed an unpaired two-sided t-test for statistical assessment of mean difference between these two categories after fisher’s z transformation of correlation r values.

### Differential gene expression analysis

We identified differentially expressed genes between progenitors and neurons by using vst normalized expression values corrected for sorter with limma R package ^72^. We retained the genes if at least 10 counts of the gene were present in more than 5% of the samples from either one of the cell type. To perform a paired differential gene expression analysis which inherently controls for donor related differences, we established the following design matrix: model.matrix(∼ CellType + as.factor(DonorID) + RIN, data). Following this, we adjusted p-values for each gene via multiple test correction with the Benjamini-Hochberg procedure ^73^, and defined significant differentially expressed genes as adjusted p-value < 0.05.

### Gene Ontology analysis

We performed gene ontology enrichment analysis by using the gprofiler2 package as the R interface to the g:Profiler tools by using GO:BP database ^74^. For differentially expressed genes, after performing DGE analysis, we categorized the genes into two groups as upregulated in progenitors (logFC < −1.5 and adjusted p-value < 0.05), and upregulated in neurons (logFC > 1.5 and adjusted p-value < 0.05) (Figure 1D). For each enrichment analysis, we applied multiple correction test, and considered only pathway enrichments with adjusted p-value lower than 5% false discovery rate as statistically significant.

### Transition Mapping (TMAP)

To evaluate the transcriptomic similarity between our *in vitro* culture system and the *in vivo* brain, we performed transition mapping analysis as described in our previous work ^26, 75^. To evaluate transcriptomic similarity to cortical laminae in the developing brain, we used previously published laminar expression data from laser capture microdissected of prenatal human brain ^31^ (H376.IIIB.02. female, 16 pcw, brainspan.org). In our comparison, genes were retained which showed expression in either cell-type and were present on the array in which the *in vivo* data was acquired. We used gene symbols to find ensemblIDs, and used ensemblIDs to match with *in vitro* data. When multiple probes were present for a given gene on the array, the probe with the highest expression per gene was used. We quantile normalized the gene expression, and we performed *in vivo* differential gene expression via limma between every two laminae. Similarly, we performed differential expression analysis in our *in vitro* cultures as described above. We applied transition mapping via RRHO2 R package with “stratified approach” to avoid misinterpretation of the discordant overlaps ^76^. In this algorithm, firstly genes were ranked based on their degree of differential expression (DDE) (i.e., −log10(p-value) × signed effect size) separately for *in vivo* and *in vitro* data. Following ranking, a hypergeometric test was applied to assess enrichment for each overlap between two datasets for a series of arbitrary cutoffs set through the highest degree to the lowest. By employing a stratified algorithm, we computed the degree of overlap. Finally, we visualized hypergeometric test −log10(p-values) as a heatmap (Figure S1G).

### Cell type specific local eQTL mapping

To perform local eQTL analysis, we conducted an association test between gene expression (retaining genes if at least 10 counts of the gene were present in more than 5% of the samples of that cell type, resulting in 24,767 and 27,638 genes for progenitors and neurons, respectively) with genetic variants within ± 1 Mb window of gene TSS for both autosomal chromosomes and X chromosome, for progenitors and neurons separately. Each gene TSS was defined as the transcription start site of the gene isoform with most upstream exon based on GTF file Homo_sapiens.GRCh38.92 (http://ftp.ensembl.org/pub/release-92/gtf/homo_sapiens/Homo_sapiens.GRCh38.92.gtf.gz).

We removed variants of low allele frequency in order to prevent one donor from strongly influencing association results. For variant selection, PLINK v1.90b3 software function was implemented to obtain donor counts per genotype group for each variant. We included only variants with at least 2 heterozygous donors and no homozygous minor allele donors, or at least 2 minor allele homozygous donors for autosomal chromosomes, and for X chromosome we retained the variants with at least 2 haploid allele counts in addition to this criteria.

For eQTL mapping, we established a linear mixed effects regression model to control for population stratification and cryptic relatedness with EMMAX software ^77^. To compute the kinship matrix, we implemented algorithm creating the identity by state (IBS) kinship matrix by excluding all genetic variants located on the same chromosome as the tested variant from non-imputed genotype data for each single variant association test (MLMe method; see ^78^. We used additional ancestry control by including the first 10 MDS components from genotype data ^79^. In order to control for unmeasured technical variables impacting gene expression, we sequentially added gene expression PCs and re-ran the genetic associations via EMMAX. For neurons, we included a covariate for FACS sorter for each run given its strong impact on gene expression.

The full association model for neurons was:

expression ∼ SNP + 10 MDS of global genotype + kinship matrix + FACS sorter + PCs of global gene expression

The full model for progenitors was:

expression ∼ SNP + 10 MDS of global genotype + kinship matrix + PCs of global gene expression

For each run, we adjusted nominal values of all gene variant associations, and defined significant associations with nominal p-value lower than 5% false discovery rate (FDR)^73^. We found that 10 PCs and 12 PCs of gene expression resulted in a maximum number of eGenes discovery in progenitors and neurons, respectively (Figure S2E). Our final eQTL model was:

Neuron:

expression ∼ SNP + 10 MDS of global genotype + kinship matrix + FACS sorter + 12 PCs of global gene expression

Progenitors:

expression ∼ SNP + 10 MDS of global genotype + kinship matrix + 10 PCs of global gene expression

In order to stringently control our association results for both number of variants and genes tested, we further implemented a hierarchical correction procedure called eigenMT-BH ^35^. Using this method, as Step 1, we adjusted the nominal p-values of the all cis SNPs separately for each gene to compute locally adjusted p-values with eigenMT method ^34^ In Step 2, locally adjusted minimum p-values for all genes were then subjected to BH procedure to obtain globally adjusted p-values. In Step 3, we defined eGenes as genes with globally adjusted p-value lower than 0.05. Then, to find other independent SNPs for those eGenes, we set the significance threshold as the maximum nominal p-value from step 1 that had corresponding globally adjusted p-value lower than 0.05.

We performed conditional analysis by using this threshold p-value gathered from the eigenMT-BH multiple correction method to identify independent significant eQTLs. To identify conditionally independent eQTLs, for each eGene (a gene significantly associated with at least one variant), we iteratively included the hard call genotype of the variant with strongest association with eGene as a covariate, and re-ran the regression model specified above (Figure S2B). We defined a variant as “conditionally independent” from the variant conditioned on, if the association of the variant with the eGene was still significant based on the initial threshold p-value. Then, we conditioned on those variants that met threshold p-value condition at the first round plus the primary variant, and identified third conditionally independent eQTLs. We applied this procedure iteratively until no additional significant eQTLs remained ^80, 81^.

### Comparison of degree of controlling population structure between EMMAX vs FastQTL

We applied FastQTL ^82^ method in nominal pass mode for different models (1) without controlling for either for population structure or technical confounders (2) controlling for only technical confounders, (3) controlling for 10 MDS of global genotype and global gene expression PCs. Following this analysis, we compared genomic inflation factors (λ_GC_) across those three groups to our data where we controlled for 10 MDS of global genotype and global gene expression, as well as the cryptic relatedness with kinship matrix.

### Bulk Fetal brain eQTL mapping

We utilized bulk fetal cortical wall eQTL data described previously ^19^. We re-analyzed data in this study with the following modifications to harmonize with the eQTL approach implemented in this study: (1) we controlled for population stratification using a linear mixed effects model as described above, and (2) we included 23 additional donors which were genotyped after the publication of the previous manuscript. We used rRNA-depleted RNA-seq data from flash frozen human fetal brain cortical wall tissues derived from 240 donors at 14-21 gestation weeks (inferred to be 12-19 post conception weeks). We excluded 4 donors for sample swap and contamination based on verifyBAMID analysis, and one donor with sex ambiguity, resulted in 235 unique donors for eQTL analysis (35 of unique donors shared with cell type specific data). Gene based annotations of the genome were derived from Homo sapiens gene ensembl version 92 (GRCh38) for eQTLs. We included only genes with at least 10 counts in 5% of donors. We normalized the data with the VST method to be used as phenotype in eQTL analysis. We also extracted genomic DNA from the same donors, and performed genotyping on a dense array (Illumina Omni 2.5+Exome) and imputation to a common reference panel (1000 Genomes Phase 3; described above). Variants were retained in the analysis if there were at least 2 heterozygous donors and no homozygous minor allele donors, or if there were at least 2 minor allele homozygous donors as for cell type specific eQTLs, as described above.

We performed local eQTL analysis to test the association between each gene’s expression and variants within the ±1 Mb window of the transcription start site of each gene. We applied linear mixed model association software, EMMAX ^77^ to control for population stratification and cryptic relatedness (as described above for cell type specific eQTL analysis). We used the linear mixed effects regression model testing association between expression of each gene and nearby genetic variants, controlling for 10 MDS genotype components, 10 PCs of gene expression, and a kinship matrix as random effect excluding the chromosome genotypes testing with the MLMe approach ^78^. After association, nominal p-values were corrected for hierarchical multiple testing using the eigenMT-BH method as described above, and we obtained independent eQTLs performing conditional analysis as described for cell type specific eQTLs above.

### Enrichment of eQTLs within functional genomic annotations

To identify enrichment of eQTLs and sQTLs within functionally annotated genomic regions, we implemented GARFIELD software to control for the distance to TSS, LD and minor allele frequency (MAF) of QTLs ^83^. We used functional genomic annotations from 25 chromatin states given in the ChromHMM BED files of Roadmap Epigenomics project from human male fetal brain^36, 84^ lifted over from hg19 to hg38, splice sites, introns, 5’ and 3’ UTR. We extracted 5’ and 3’ splice sites (splice donor and splice acceptor sites) from Homo_sapiens.GRCh38.92 GTF file implementing hisat2_extract_splice_sites.py algorithm of hisat2 software (Kim, Langmead, and Salzberg 2015), and defined exon-intron boundaries as the region between those splice sites. 5’ and 3’ UTR were also defined based on the coordinates in the same GTF file. For all e/sQTLs, we extracted the p-value from the strongest association for each variant (with minimum p-value) in the case that one variant was associated with multiple genes/intron junctions. To create annotations files, we considered a variant overlapping with a functional element if the variant itself or any of the variants in high LD within 500kb (r^2^ > 0.8) overlapped with each of annotation categories. LD pruning^83^ was performed at r^2^ > 0.01 within GARFIELD software. Following this, a logistic regression model controlling for the distance to TSS, LD proxies and MAF binned for five quantiles was performed with GARFIELD software for enrichment at eigenMT-BH p-value thresholds defined in eQTL analysis. The effective number of annotations were estimated and multiple testing adjusted p-values were computed by the software to identify enrichment of eQTLs within defined annotations.

### RNA binding protein motif analysis

We performed sQTL enrichment in RNA binding protein binding sites via GARFIELD as described above. In this analysis, we used BED files including RNA binding protein sites from a CLIP-seq database as annotation files ^44^, and assessed significant enrichment of cell type specific sQTLs for binding sites of each RBP.

### Allele specific expression analysis pipeline

To identify sites with Allele Specific Expression (ASE), we applied the ASEReadCounter algorithm from GATK tools ^85^ to the RNA-seq data remapped with WASP to reduce mapping bias and to discard duplicate reads. For each donor, we counted allele specific reads overlapping with bi-allelic variants identified in the genotypeVCF files. We retained only variants with at least 5 heterozygous donors, and at least 10 counts from either allele. ASE can be falsely called when genotyping errors are present in the dataset. We used two approaches to identify and remove potential genotyping errors: (1) We detected wrongly called variant genotypes by assessing concordance between genotypes called by DNA versus RNA ^37^. We removed variants that were called homozygous based on the genotype data when at least 10 counts of the alternate allele were present in the RNA-seq data, (2) we discarded variants where at least 7 heterozygous donors based on genotype data have zero counts for one of the alleles, which may indicate a donor falsely called as heterozygote when in truth the donor is a homozygote (given that (1/2)^7 = 0.008, meaning that probability of having all donors receiving an imprinted allele from either mother or father is low). Because ASEReadCount does not disambiguate the strandedness of reads, it is not possible to confidently assign reads overlapping with multiple gene annotations to a specific gene ^64^. Therefore, if a variant overlapped with more than one gene annotation, we removed the variant by implementing findOverlaps function from IRanges R package ^86^ for genes based on their genomic coordinates defined GTF file Homo_sapiens.GRCh38.92 (http://ftp.ensembl.org/pub/release-92/gtf/homo_sapiens/Homo_sapiens.GRCh38.92.gtf.gz).

To evaluate allelic imbalance, we used DESeq2 with the design: design = ∼0 + RNAid + Allele. Excluding homozygous donors, we computed the log2 fold change of non-reference allele counts over reference allele counts and used a Wald test to detect allelic imbalance. Multiple test correction was performed with the Benjamini and Hochberg method, and we defined significant ASE sites as those with adjusted p-values lower than 0.05.

To compare eQTLs with ASE sites (Figure S3E-S3H), we extracted eQTLs associations with the same filtering criteria of variants used for ASE analysis (at least 5 heterozygous donors and overlapping with at least 10 RNA-seq reads). We also extracted eGenes (defined based on significant eigenMT-BH global p-value) with at least 10 counts per donor.

### Quantification of Intron Excisions

To identify alternatively excised introns, separately for each cell type, we extracted exon-exon junctions from WASP-mapped RNA-seq data in BAM format via regtools function where reads map to a minimum of 6 nt of each exon ^87^. Next, we processed those junctions that are called *intron excisions* or *exon-exon junctions* with the pipeline provided by Leafcutter software ^40^. Firstly, intron excisions with shared splice junctions were clustered together applying an iterative procedure until each cluster has at least 50 reads across donors and introns with maximum 50 kb length, separately for progenitors and neurons. For differential splicing analysis, we performed clustering by combining exon-exon junctions files from each cell type. For each cluster, intron excisions supported by at least one count in more than 5 donors (within each set of donors contributing to the 3 different sQTL analyses for that cell type (progenitor, neuron) or tissue class (fetal brain bulk); or for differential splicing analysis across donors from both cell types used (progenitor + neuron) were retained. We further calculated intron excision ratios, and filtered out introns represented in less than 40% of donors (within each set of donors contributing to the 3 different sQTL analyses for that cell type (progenitor, neuron) or tissue class (fetal brain bulk); or for differential splicing analysis across donors from both cell types used (progenitor + neuron) with algorithm. We referred to each intron excision ratio as *percent spliced in* (PSI) that corresponds to the usage of each intron compared to other introns in the same cluster. Standardized and quantile normalized intron excision ratios, and global alternative splicing PCs computed with those ratios were used for downstream analysis.

### Differential splicing analysis

To perform differential splicing analysis, we used quantile normalized PSI values as input to the limma package ^72^. Identical to differential expression analysis, neuron splice ratios were corrected for batch including FACS machine used for sorting with function. Batch corrected neuron splice ratios were combined with progenitor data. We implemented a paired differential splicing analysis inherently controlling donor related differences with the design matrix: model.matrix(∼CellType + as.factor(DonorID) + RIN, data). We defined intron junctions with adjusted p-values from via multiple test correction with Benjamini-Hochberg procedure ^73^ lower than 0.05 as significant differentially spliced introns.

### Splicing QTL mapping

We performed cell type specific splicing QTL analysis by testing the association of PSI with the genetic variants located within the ± 200 kb window from starting and end points of the splice junctions for autosomal chromosomes and the X chromosome. Identical to local eQTL analysis, we used only genetic variants that met the following criteria: if there were at least 2 heterozygous donors and no homozygous minor allele donors, or if there were at least 2 minor allele homozygous donors.

We used standardized and normalized intron excision ratios (percent spliced in) calculated by leafcutter as the phenotype for sQTL mapping. EMMAX ^77^ was used to test for association between SNPs within a cis-region of ± 200kb of the intron cluster and intron ratios within cluster. We controlled for population stratification and cryptic relatedness as described above for eQTL mapping. Also, we controlled for unmeasured technical variables impacting alternative splicing by sequentially adding global splicing PCs to the genetic associations via EMMAX. Again for neurons, we additionally controlled for FACS sorter for each run given its strong impact on splicing as well.

The full model for neurons was:

PSI ∼ SNP + 10 MDS of global genotype + kinship matrix + FACS sorter + PCs of global splicing

The full model for progenitors was:

PSI ∼ SNP + 10 MDS of global genotype + kinship matrix + PCs of global splicing

For every run, we adjusted nominal values of all PSI variant associations, and defined significant associations with lower than at 5% false discovery rate (FDR) ^73^. We found that 1 PC and 1 PC across the PSI matrix resulted in a maximum number of intron excisions with at least one significant association in progenitors and neurons, respectively (Figure S2E). Our final sQTL model was: Neuron:

PSI ∼ SNP + 10 MDS of global genotype + kinship matrix + FACS sorter + 1 PCs of global splicing

Progenitors:

PSI ∼ SNP + 10 MDS of global genotype + kinship matrix + 1 PCs of global splicing

Implementing a hierarchical correction procedure called eigenMT-BH ^35^, firstly, we adjusted the p-values of the all cis SNPs strongest association separately for each intron excision to compute locally adjusted p-values with the eigenMT method ^34^, and then locally adjusted minimum p-values for all intron excisions were subjected to the BH procedure giving globally adjusted p-values. Intron excision with corresponding global p-value lower than 0.05 were considered as significant alternative splicing events. In order to find other independent significant sQTLs in addition to the ones associated with lowest p-values, we applied conditional analysis at eigenMT-BH p-value threshold as described for eQTL analysis.

For bulk fetal cortical tissue sQTL mapping, we applied the same strategy used for cell type specific sQTLs, and found the following model maximized significant intron junctions discovery:

PSI ∼ SNP + 10 MDS of global genotype + kinship matrix + 5 PCs of global splicing

After calculating eigenMT-BH threshold p-value, we performed conditional analysis to define independent significant sQTLs.

To find genes overlapping with intron excision, we annotated intron junctions by using Leafcutter based on genomic coordinates and gene model provided in GTF file Homo_sapiens.GRCh38.92. Intron junctions assigned as cryptic 5’, cryptic 3’, novel annotated pair were considered as novel splicing events for the genes overlapped with junctions. For unannotated splice sites for *AS3MT* and *ARL14EP* genes, we additionally checked for overlap with known splice sites up to Ensembl Release 100.

### Estimation of m-values for cross cell comparison

We estimated m-values to assess cell type specificity of SNP-gene or SNP-intron excision pairs with Metasoft ^88^. Prior to software implementation, we extracted e/sQTLs from the neuron data corresponding to conditionally independent progenitor e/sQTLs to see overlap of sharing significant progenitor e/sQTLs with neuron eQTLs. Similarly, we extracted e/sQTLs from the progenitor data corresponding to conditionally independent neuron e/sQTLs to see overlap of sharing significant neuron eQTLs with progenitor e/sQTLs. We estimated standard errors via dividing beta estimates from EMMAX by t-statistics for each association p-value. We considered associations shared across different QTLs for the m-value > 0.9.

Similarly, in order to find significant progenitor/neuron e/sQTLs shared with fetal bulk e/sQTLs, we extracted e/sQTLs from the fetal bulk data corresponding to conditionally independent progenitor/neuron e/sQTLs, and defined shared at m-value > 0.9.

### Comparison of cell-type specific vs fetal and adult bulk brain e/sQTLs

We considered an overlap of e/sQTLs between two datasets when the index e/sSNPs were in LD (r^2^>0.8 where LD was calculated in our sample population) and the eSNP-eGene/sSNP-intron pairs were shared. To determine the total number of eSNP-eGene/sSNP-intron pairs as the universe for enrichment analyses, we pruned all variants associated with each gene per gene for r^2^ > 0.01 by using PLINK command plink --indep-pairwise 50 5 0.01. To determine if different proportions of sharing were observed between two cell types, we performed an odds ratio test described here ^89^.

To test temporal specificity of cell type specific e/sQTL data, we downloaded GTEx data adult brain e/sQTL data ^24^. We called loci from the two datasets as colocalized when, (1) index adult brain e/sQTLs are found within LD buddies of cell type specific e/sQTLs at LD r^2^ > 0.8 (where LD is calculated using either the European population from 1000 Genomes or our study’s population), and (2) when the cell-type specific e/sQTL data conditioned on index adult brain e/sQTLs, the index brain e/sQTL no longer survives the global significance threshold.

### LD-thresholded colocalization with brain disorders and traits GWAS

To find eQTLs and sQTLs colocalized with index GWAS loci, we performed LD-thresholded colocalization analysis for each cell type separately ^47^. We used summary statistics of GWAS for schizophrenia (SCZ) ^1^, major depression disorder (MDD) ^90^, bipolar disorder (BP) ^2^, educational attainment (EA)^91^, neuroticism ^92^, IQ ^5^, cognitive performance (CP) ^91^, attention-deficit/hyperactivity disorder (ADHD) ^93^, Alzheimer’s disease (AD) ^94^, Parkinson’s disease (PD) ^95^, Insomnia ^96^, epilepsy^97^, autism spectrum disorder (ASD) ^98^, and cortical thickness and surface area from ENIGMA project ^4^. We liftovered positions of variants in GWAS summary statistics from hg19 to hg38 with function from R rtracklayer package ^99^. Variant rsids were assigned with dbSNP151 based on positions of variants in summary statistics data. To define index GWAS SNPs at genome-wide significant threshold p-value (5×10^-8^), we implemented a clumping procedure, where we defined two LD-independent GWAS signals so as to have pairwise LD r^2^ < 0.5 based on LD matrix computed with European population of 1000 Genomes (1000G European phase 3). Prior to clumping, duplicated rsIDs in 1000G EUR genotype files were assigned with unique names, and BIM files were modified for each chromosome. Following a unique id assignment, BIM files were merged back to BED and FAM files with --bmerge function of PLINK1.9 software (plink --bfile BED file --bmerge modified_BIM file). Since all GWAS we leveraged in our colocalization analysis have been conducted in populations of European ancestry, and our study population is multi-ancestry, we computed LD r^2^ separately within these two different populations. We considered the index eQTL or sQTL SNP coincident with the index GWAS SNP if the pairwise LD r^2^ between them was greater than 0.8 based on either the LD matrix computed via either European 1000 Genomes Phase 3 data or our study population. Following that, we performed a conditional eQTL/sQTL analysis by conditioning on the coincident index GWAS SNP. If the association of index QTL and gene expression or intron excision was no longer significant based on p-value thresholds defined with eigenMT-BH method for each dataset, we identified that cell type specific and fetal bulk eQTL/sQTL as a colocalized loci with the given GWAS trait. Since GTEx raw data is not available publicly, conditional analysis was not performed to infer colocalization.

### Transcription factor motif analysis

We used motif breaker R to detect the disruption of the transcription motif binding site where there was a variant within a chromatin accessibility peak (Figure 3D)^100^.

### TWAS analysis

We performed transcriptome wide association analysis for progenitor and neurons separately with FUSION software (http://gusevlab.org/projects/fusion/, ^52^). First, we obtained a set of variants shared between the genotypes from 1000 Genome European phase 3 ^68^ and our study population restricted to variants described for eQTL analysis, and removed monomorphic variants within European genotype data. We estimated cis-heritability of genes (including variants within ± 1 MB window of the TSS) and intron junctions (including variants within ± 200kb window of two ends of intron junctions) with GCTA software ^101^ by controlling for same covariates for global gene expression/splicing and 10 PCs of global genotypes used in e/sQTL analysis. VST normalized gene expressions were further subject to quantile normalization for heritability estimation. 1,703/973 genes and 6,728/6,799 intron junctions were significantly cis-heritable in progenitors/neurons for heritability p-value < 0.01. To determine the method to be used to estimate genetic component of gene expression/splicing (weights), we performed leave-one-out cross validation ^102^ for the prediction models including LASSO regression ^103^, Elastic-net regression ^104^ and EMMAX ^77^ within FUSION software. We used the weights computed from the prediction model with the highest cross validation R^2^ (the highest performance) per gene/intron junction for downstream analysis for progenitor, neuron and fetal bulk brain tissue. For adult brain bulk tissue data, we obtained the weights of genes and intron junctions from CommonMind Consortium study ^53^.

Before running TWAS analysis, we prepared GWAS summary statistics for schizophrenia (SCZ)^1^, major depression disorder (MDD) ^90^, educational attainment (EA)^91^, neuroticism ^92^, IQ ^105^, Alzheimer’s disease (AD) ^106^, Parkinson’s disease (PD) ^95^, and Global Surface Area (GSA) and average thickness from ENIGMA study ^4^ with following adaptations: (1) we obtained common variants found both in genotype files from our study and in GWAS summary statistics; (2) we calculated z-score by dividing the beta coefficient by the standard error if the beta coefficient was available in the summary statistics, or dividing the natural logarithm of odds ratio by the standard error if odds ratio was given in the summary statistics; (3) The sign of the z-score were matched based on the allelic directionality of weights from FUSION software.

To perform TWAS analysis, we tested the association between the predicted gene expression/splicing (w) and brain traits listed above (Z) by implementing the algorithm *Z_TWAS_= w’ Z/sqrt(w’Dw)* where D is the LD matrix as the covariance among all cis-variants from the FUSION software ^52, 53^. Since the population structure of our dataset was different from European neuropsychiatric GWAS, we performed TWAS analysis separately with different LD estimates computed based on our study or European population from 1000 Genomes Phase 3 as the covariance. For variants missing in GWAS summary statistics which existed in our study’s genotypes, we implemented IMPG imputation ^107^ allowing 40% of missing variants as maximum ratio within the FUSION algorithm.

To identify genes/intron junctions not driven by co-expression, we defined jointly independent genes/intron junctions through performing summary-statistic-based joint analysis ^108^, where we replaced SNPs with genes/intron junctions as described in previous work ^53^ within the FUSION software. Implementing genes/intron junctions to the model one at a time in decreasing order of significance, we evaluated whether the conditional TWAS test remained significant. Those with significant conditional TWAS association were defined as jointly independent.

